# Activity Profiling of Mycobacterial L,D-Transpeptidases

**DOI:** 10.64898/2026.02.05.698683

**Authors:** Karl L. Ocius, Rachel E. Sanborn, Ananya Naick, Leighanne A. Brammer Basta, Marcos M. Pires

## Abstract

Antimicrobial resistance poses major therapeutic challenges, particularly for multidrug-resistant mycobacterial infections caused by *Mycobacterium tuberculosis* (*Mtb*) and non-tuberculous mycobacteria (NTM). L,D-Transpeptidases (Ldts) are attractive drug targets due to their essential role in peptidoglycan cell wall crosslinking, yet existing assays suffer from low throughput and limited sensitivity. We report a versatile, bead-based platform for high-throughput analysis of Ldt activity and inhibitor discovery. We incubated peptidoglycan stem peptides, either naturally harvested or synthetically immobilized on abiotic surfaces, with Ldts and a fluorescent acyl acceptor to quantitatively monitor crosslinking. After optimizing assay parameters, we profiled six Mycobacterium smegmatis Ldt paralogs, including the first characterization of a class 6 Ldt with chemically defined substrate sequences. Utilizing a series of acyl acceptors, we demonstrated modifications within the acyl acceptor that are tolerated by mycobacterial Ldts. Screening of β-lactam antibiotics revealed potent inhibition by (carba)penems, while cephalosporins, monobactams and penams showed negligible activity. The assay achieved excellent performance metrics and was successfully adapted to ELISA and 96-well formats, providing a powerful tool for discovering Ldt-targeted therapeutics against tuberculosis and related infections.

## INTRODUCTION

Tuberculosis (TB), caused by the bacterium *Mycobacterium tuberculosis* (*Mtb*), remains one of the world’s deadliest infectious diseases, claiming 1.23 million lives in 2024 alone.^1^ The disease burden is unevenly distributed across patient populations, with TB representing the leading cause of mortality among people living with HIV.^2, 3^ The emergence of multidrug-resistant (MDR) and extensively drug-resistant (XDR) strains continues to erode therapeutic options,^1, 4^ while infections caused by non-tuberculous mycobacteria (NTM) are rising worldwide.^5^ Together, these trends underscore an urgent need for novel antimycobacterial agents. Yet antibiotic development has stagnated, and the broader antimicrobial resistance crisis is projected to cause 10 million deaths annually by 2050.^6^ Addressing this challenge requires not only new drug candidates but also innovative tools to identify and validate them.

The bacterial cell wall, specifically its peptidoglycan (PG) layer, has long served as a validated therapeutic target. This mesh-like structure provides mechanical strength and defines cellular shape, making it essential for bacterial survival.^7, 8^ In mycobacteria, PG architecture has glycan strands composed of alternating *N*-acetylglucosamine and *N*-acetylmuramic acid that are crosslinked by peptide bridges, creating a complex and inherently heterogenous macromolecular network. A prominent feature of PG is the high degree of crosslinking observed between neighboring peptide strands. Two enzyme families catalyze these crosslinks: the D,D-transpeptidases (penicillin-binding proteins, PBPs) and the L,D-transpeptidases (Ldts) (**Fig. 1a**).^9^ Unlike PBPs, which utilize a serine-based catalytic mechanism to form 4→3 peptide crosslinks, Ldts employ a cysteine-based mechanism to generate 3→3 crosslinks between adjacent peptide stems (**Fig. 1b**). The first step in Ldt-catalyzed crosslinking involves acylation of the active site cysteine by the carbonyl of the acyl donor substrate (donor strand), forming a covalent acyl-enzyme intermediate, and releasing the terminal group D-Alanine (D-Ala). At this intermediate stage, there is a thioester intermediate between the Ldt and the truncated donor strand. Next, the ε-amino group of *meso*-diaminopimelic acid (*m*-DAP) from an adjacent acceptor peptide strand attacks this activated acyl intermediate, generating the crosslink and regenerating free Ldt enzyme.^10^

**Figure 1.**
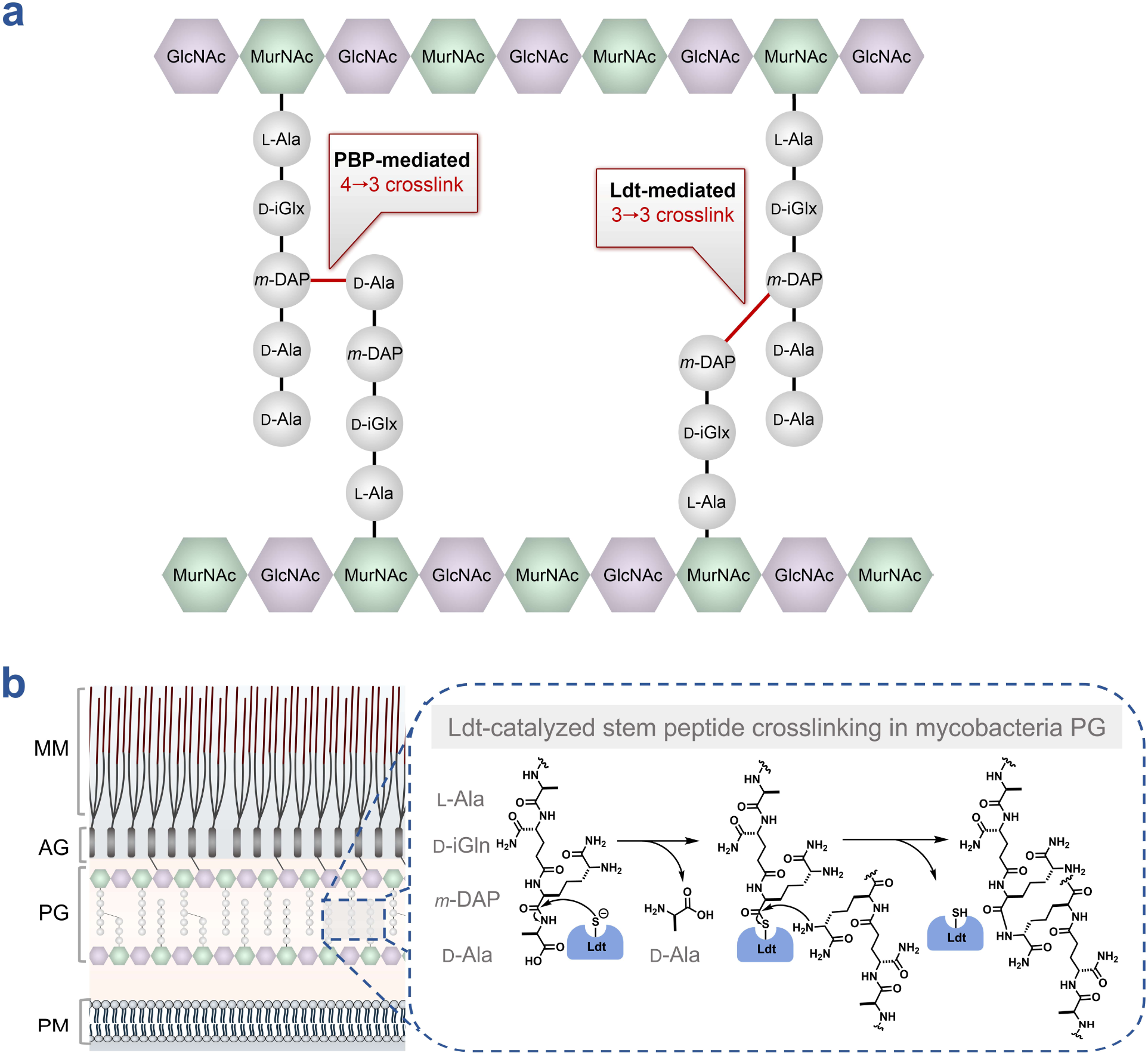
Composition and stem peptide crosslinking within the cell wall of mycobacteria. (**a**) PG consists of glycan strands composed of alternating *N*-acetylglucosamine (GlcNAc) and *N*-acetylmuramic acid (MurNAc) to which a stem peptide is covalently attached. Adjacent stem peptides are crosslinked to form a macromolecular network. Crosslinking is catalyzed by two enzyme families: D, D-transpeptidases (penicillin-binding proteins, PBPs) which generate 4→3 crosslinks, and the L, D-transpeptidases (Ldts), which generate 3→3 crosslinks. (**b**) The mycobacteria cell envelope comprises the mycomembrane (MM), arabinogalactan (AG), peptidoglycan (PG), and plasma membrane (PM). Ldts catalyze stem peptide crosslinking within the periplasm via a two-step mechanism. The catalytic cysteine first performs a nucleophilic attack on a donor stem peptide, forming an enzyme–peptide intermediate and releasing the *C*-terminal D-alanine (D-Ala). The ε-amino group of the *meso*-diaminopimelic acid (*m*-DAP) residue of an adjacent acceptor stem peptide then attacks this intermediate, resulting in formation of a covalent 3→3 crosslink and regeneration of the free enzyme.

While mycobacteria encode both enzyme classes, Ldts predominantly mediate PG crosslinking in these species and are essential for cell wall integrity, making them critical yet challenging therapeutic targets. Genetic deletion studies have confirmed that Ldts are essential for mycobacterial survival, with their absence severely compromising cell wall stability and restricting bacterial growth.^11–15^ These phenotypic observations establish Ldts as validated therapeutic targets whose inhibition could effectively expand the arsenal of antimycobacterial agents to combat TB infections. However, Ldts are notoriously difficult to inhibit due to their unique active site architecture, which restricts β-lactam access and often results in unstable or readily reversible covalent adducts.^16–20^ This challenge is compounded by chromosomally encoded β-lactamases in *Mtb* and NTM species that hydrolyze and inactivate many β-lactams before they reach their targets,^21–24^ leaving only carbapenems and penems as effective Ldt inhibitors. While recent clinical success with β-lactam/β-lactamase inhibitor combinations has renewed interest,^25–28^ systematic discovery of novel Ldt inhibitors remains hampered by significant methodological limitations.

A major obstacle to Ldt-targeted drug discovery has been the difficulty in characterizing enzyme activity, which requires synthesis of peptidyl substrates containing *m*-DAP, a building block that remains challenging to produce using standard solid-phase peptide synthesis.^29^ Compounding this challenge, existing assays are insufficient to accurately monitor the two-step transpeptidation reaction and characterize the 3→3 crosslinked products formed during catalysis, which are often polymeric and heterogenous in nature.^12, 30–32^ There have been prior efforts guided towards the analysis of broad inhibition of Ldts, which have attempted to circumvent these challenges. Thiol-reactive colorimetric and fluorogenic probes have been used to indirectly monitor inhibition by assessing overall cysteine reactivity; however, these approaches provide little insight into features unique to Ldts.^30, 33^ Similarly, D-alanine oxidase (DAAO) has been repurposed to detect the release of D-Ala that accompanies the first step of the transpeptidation reaction.^34^ More recently, a rotor-fluorogenic D-amino acid probe was shown to report on changes in bond rotation upon crosslinking, enabling detection of both Ldt and PBP activity.^35^ Finally, work from the Beatty and Carlson groups has revealed the range of detectable transpeptidase targets using fluorescently labeled β-lactams combined with gel-based imaging.^36–41^

Given the possible redundancy of Ldts in many organisms, including mycobacteria which encode ∼5-6 individual Ldts, it remains to be elucidated how individual transpeptidases (including PBPs) operate within the inherent heterogeneity of PG matrices. Some studies suggest, at least in terms of specific PG labels, that the transpeptidation with analogs of the stem peptide could be specific to a single PBP in *S. aureus*^42^ and a single Ldt in *E. coli*.^43^ We hypothesized that specific PG strand structures temporally and spatially coordinate Ldt activity. The extent of this structural variation is substantial; a single bacterial strain can display over 250 unique muropeptides.^44^ We, and others, recently demonstrated that amidation of *iso*-glutamic acid with the muropeptide is critical for transpeptidase processing in live cells.^45–48^ Given this vast range of unique PG structures within a single cell, Ldt processing could be potentially likely driven, to varying extents, by muropeptide primary sequence.

Existing assays for evaluating Ldt inhibition and reactivity with covalent modifiers rarely incorporate substrates that reflect the structural complexity of native PG, limiting mechanistic insight into Ldt function. To address this gap, we developed a high-throughput transpeptidation platform that reconstitutes and quantifies both steps of Ldt catalysis. A synthetic acyl-donor peptide is immobilized on solid supports (or native sacculi), and enzyme-mediated incorporation of a fluorescent acyl-acceptor is quantified by flow cytometry. This configuration captures the full catalytic cycle (acylation of the enzyme by the bead-bound donor followed by nucleophilic resolution by the fluorescent acceptor) yielding a crosslinked product whose formation is directly reported by fluorescence levels. Active-site Ldt inhibitors block crosslinking and produce a proportional loss of signal. Applying this system, we delineated transpeptidase activities and acyl acceptor processing across multiple Ldt paralogs and assessed inhibition by a focused β-lactam library. The compatibility of the assay with both flow-cytometric and ELISA-based detection expands its utility, and its robust performance metrics underscore its value for mechanistic studies and screening applications.

## RESULTS AND DISCUSSION

Several pathogenic bacteria, including mycobacteria, incorporate the unusual amino acid *m*-DAP at the third position of the PG muropeptide (**Fig. 2a**).^49, 50^ The inherent symmetry of this building block presents a significant synthetic challenge. As a result, *m*-DAP is not commercially available, and existing synthetic routes are lengthy and proceed in low overall yield.^51–56^ These difficulties in accessing *m*-DAP have greatly hampered the development of chemical probes designed to study *m*-DAP-containing structures and pathways. To address these challenges, we and others have developed stem peptide-mimicking chemical probes, including variants employing structural surrogates such as *meso*-cystine (*m*-CYT) and *meso*-selenolanthionine (*m*-SeLAN) in place of *m*-DAP (**Fig. 2a**).^42, 45, 57–62^ These analogues are well tolerated across multiple bacterial species, including mycobacteria, and have since enabled interrogation of Ldt activity within the mycobacterial cell wall in live cells.^57, 58^

**Figure 2.**
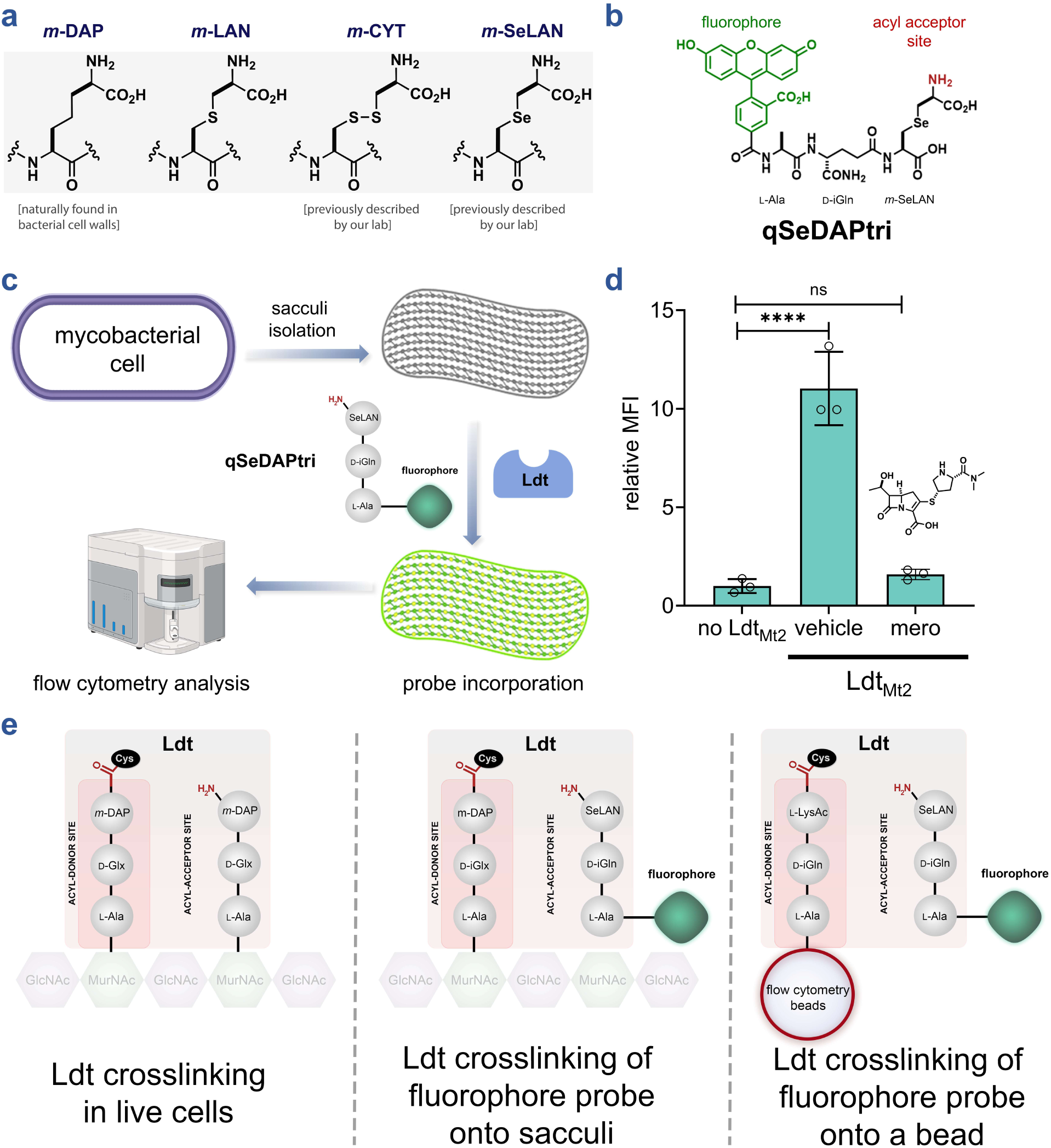
Assessment of Ldt-catalyzed crosslinking of fluorescent acyl acceptor onto M. smegmatis sacculi. (**a**) Schematic representation of *m*- Dap and the analogues incorporated within the PG of mycobacteria. (**b**) Schematic of the fluorescent tripeptide acyl acceptor **qSeDAPtri.** (**c**) PG sacculi is isolated from other cellular components of *M. smegmatis*. Once isolated, incubation of **qSeDAPtri** and the purified Ldt result in a fluorescent sacculi that can be analyzed by flow cytometry. (**d**) Flow cytometry analysis of isolated PG sacculi incubated with 20 µM **qSeDAPtri** in the presence, and the absence of 2 µM Ldt_Mt2_ for 2 h. Mean fluorescence intensity (MFI) represents the ratio of fluorescence relative to the control (no Ldt_Mt2_), quantified from 10000 events. Statistical significance was determined by one-way ANOVA (ns = not significant, **** p<0.0001). (**e**) Illustrated overview depicting stem peptide processing within bacterial PG in vivo, incorporation of fluorescent stem peptide mimics into isolated PG sacculi, and probe crosslinking to a single acyl acceptor stem peptide mimic immobilized on an abiotic surface.

We leaned on these prior efforts to more faithfully mimic native Ldt substrates, including the synthesis of a fluorescent tripeptide acyl acceptor strand (**qSeDAPtri**, **Fig. 2b**). Restricting the substrate to a tripeptide ensured that it functioned exclusively as an acyl acceptor, as it lacks the terminal D-Ala required to serve as an acyl donor during the crosslinking reaction. In addition, we included a fluorophore onto the *N*-terminus of the probe, which is a site within muropeptide probes that we previously showed tolerate modification and still be processed by Ldts.^58, 63^ To test **qSeDAPtri** processing by Ldts, we initially reasoned that isolated sacculi could serve as the acyl donor chains (**Fig. 2c**). Importantly, the use of sacculi, as a highly diverse and comprehensive set of acyl donors, can potentially offer unique insights into the breadth of substrates that can operate with Ldts. Incorporation of **qSeDAPtri** *via* Ldt mediated crosslinking was expected to result in higher levels of sacculi-associated fluorescence; this step can readily be monitored by flow cytometry (SaccuFlow), a platform we recently disclosed to broadly enable higher-throughput detection of structural changes in isolated sacculi from various species of bacteria.^64, 65^

As a proof of principle for sacculi-based analysis, we focused on Ldt_Mt2_, the best characterized Ldt in *Mtb*. It is well established that Ldt_Mt2_ is responsible for the majority of 3→3 crosslinks in the *Mtb* PG, is essential for maintaining cell wall integrity during growth and stress and represents an important target for antibacterial drug development.^11–13, 66–70^ To benchmark our system, Ldt_Mt2_ was incubated with sacculi and **qSeDAPtri** and the resulting sacculi-associated fluorescence was monitored by flow cytometry. Our results showed that incubation of isolated mycobacterial sacculi with **qSeDAPtri** and Ldt_Mt2_ yielded a marked fluorescence increase over sacculi treated with the probe in the absence of the Ldt enzyme, confirming assay feasibility (**Fig. 2d**). To confirm that fluorescence resulted from Ldt_Mt2_ enzymatic activity, we pre-treated the enzyme with meropenem. Meropenem has been previously shown to block processing by Ldts by covalent trapping of the active site cysteine.^18, 68, 71^ In our platform, meropenem treatment reduced sacculi-associated fluorescence to near background levels, demonstrating that this approach enables high-throughput discovery of novel Ldt inhibitors. In principle, this strategy can also be adapted to probe species- and strain-specific acyl-donor preferences through sacculi isolation.

While sacculi provide several advantages for probing Ldt activity (e.g. accessibility, native acyl-strand composition, facile isolation by standard laboratory techniques), the structural heterogeneity of acyl-donor substrates (present as muropeptides within the sacculus) limits the ability to conduct rigorous structure–activity relationship analyses of inhibitors and substrates (**Fig. 2e**). To this end, muropeptides, although initially biosynthesized as canonical pentapeptides of uniform composition, subsequently undergo extensive structural modifications that include removal of several amino acids (generating tetrameric, trimeric, dimeric, and denuded strands), amidation of D-*i*Glu residues, and incorporation of non-canonical amino acids in the 4^th^ and 5^th^ position.^7, 72^ Additionally, PG-associated molecules, including lipoteichoic acids and wall teichoic acids in Gram-positive bacteria, or the mycolyl-arabinogalactan layer in mycobacteria, must be removed to ensure enzyme access.^73–79^ Thus, isolating muropeptides reflective of mycobacterial PG diversity are not only time-consuming but also introduce batch-to-batch variability, ultimately limiting reproducibility and throughput.^75, 77, 80^ These limitations prompted us to develop an alternative strategy. In this iteration, a synthetic acyl donor peptide is immobilized onto polystyrene beads, and Ldt-mediated crosslinking of a fluorescent acyl acceptor is quantified by flow cytometry. Critically, both acyl donors and acyl acceptor strands are synthetic with a precisely defined structure. This approach recapitulates both catalytic steps: enzyme acylation by the bead-bound donor (with D-Ala release) and subsequent crosslink formation with the fluorescent acceptor, rendering beads fluorescent.

To implement this bead-based strategy, we synthesized an acyl donor peptide bearing an *N*-terminal propargyl group (**tetAcK-yne**, **Fig. 3a**). The alkyl group was used to covalently link the synthetic acyl donor strand to azide-functionalized polystyrene beads *via* copper-catalyzed azide-alkyne cycloaddition (CuAAC). To ensure **tetAcK-yne** functioned exclusively as the acyl donor, we acetylated the lysine ε-amino group, thereby preventing it from acting as an acceptor.^58, 63^ We first validated this chemical step by performing CuAAC with fluorescein-alkyne onto the azide-bearing beads and analyzing the fluorescence levels of the beads by flow cytometry (**Fig. S1**). Upon the addition of copper reagents, there was a marked increase in bead-associated fluorescence. After confirming successful bead functionalization *via* click chemistry, we conducted a series of experiments to optimize transpeptidation reactions using the bead-based reporter system. As with the sacculi experiments, **qSeDAPtri** again served as the fluorescent acyl acceptor substrate for Ldt_Mt2_-mediated crosslink analysis. In the absence of the enzyme, bead fluorescence levels were low, suggesting minimal non-specific binding of **qSeDAPtri** onto the bead support (**Fig. 3b**). Similarly, the incubation of the beads with Ldt_Mt2_ but in the absence of **tetAcK-yne** resulted in similarly low levels of bead fluorescence, which indicates that the fluorescence must originate from crosslink formation with the acylated strand. Satisfyingly, **tetAcK-yne**-linked beads incubated with both the acyl acceptor strand and the enzyme resulted in a significant increase in bead associated fluorescence. Upon the pre-treatment of Ldt_Mt2_ with the β-lactam meropenem, bead fluorescence returned to background levels. Together, these results confirmed that our platform specifically reports Ldt-catalyzed transpeptidation.

**Figure 3.**
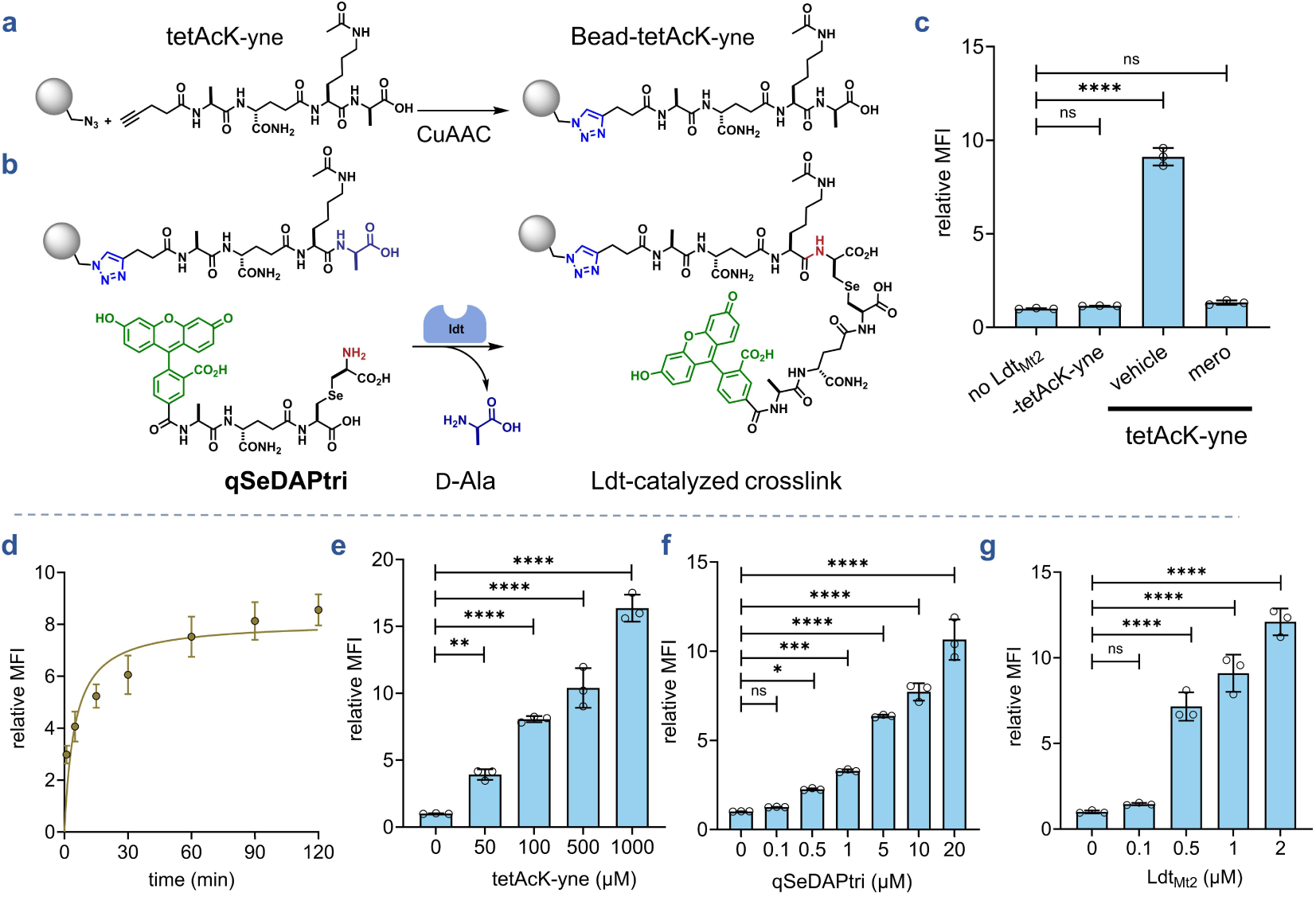
(**a-b**) Schematic representation of the L,D-transpeptidation assay. A fluorescent acyl-donor PG stem-peptide mimic (**tetAcK-yne**) was immobilized onto flow-cytometry–compatible polystyrene beads via copper-catalyzed azide–alkyne cycloaddition (CuAAC). Incubation of bead-bound **tetAcK-yne** with a fluorescent acyl-acceptor peptide (**qSeDAPtri**) and an Ldt results in enzyme-mediated crosslinking of the fluorescent peptide onto the bead surface with loss of the terminal D-alanine residue from **tetAcK-yne** . Fluorescence, reflecting Ldt activity, will be quantified by flow cytometry. (**c**) Flow cytometry analysis of beads bearing 100 µM **tetAcK-yne** incubated with 20 µM **qSeDAPtri** in the presence, absence, or inhibition of 2 µM Ldt_Mt2_ by meropenem for 2 h. (**d**) Time course of crosslinking. Beads bearing 100 µM **tetAcK-yne** were incubated with 20 µM **qSeDAPtri** and 2 µM Ldt_Mt2_ over 120-min intervals. (**e**) Concentration scan of acyl-donor loaded on beads. Beads bearing 50, 100, 500, or 1000 µM **tetAcK-yne** were incubated with 20 µM **qSeDAPtri** and 2 µM Ldt_Mt2_ for 1 h. (**f**) Concentration scan of acyl-acceptor substrate. Beads bearing 100 µM **tetAcK-yne** were incubated with 0.1, 0.5, 1, 5, 10, or 20 µM **qSeDAPtri** and 2 µM Ldt_Mt2_ for 1 h. (**g**) Enzyme-concentration scan. Beads bearing 100 µM **tetAcK-yne** were incubated with 5 µM **qSeDAPtri** and 0.1, 0.5, 1, or 2 µM Ldt_Mt2_ for 1 h. Mean fluorescence intensity (MFI) represents the ratio of fluorescence relative to the control (no Ldt_Mt2_), quantified from 10000 events. Statistical significance was determined by one-way ANOVA (ns = not significant, **** p<0.0001).

### Systematic optimization of assay parameters

Having established proof-of-concept, we systematically profiled reaction conditions to maximize assay sensitivity and the dynamic range. We first determined optimal reaction time by monitoring Ldt_Mt2_ activity over time. Reactions were quenched with PBS containing trifluoroacetic acid, which protonates the catalytic cysteine to prevent further catalysis. For all assays, upon the completion of the assay period, beads were washed with Tween-20 containing buffer to remove non-covalently associated **qSeDAPtri**. Time-course analysis revealed fluorescence plateaued after approximately 1 h (**Fig. 3c**), which we adopted for subsequent experiments. We next probed how the **tetAcK-yne** concentration could modulate the dynamic range. Azide-functionalized beads underwent CuAAC with varying **tetAcK-yne** concentrations, followed by incubation with the Ldt enzyme and **qSeDAPtri**. Our data showed that 100 µM of **tetAcK-yne** yielded optimal signal levels (**Fig. 3d**), balancing substrate availability with background minimization. **qSeDAPtri** concentration was probed by incubating **tetAcK-yne** beads with Ldt_Mt2_ and varying acyl acceptor concentrations (**Fig. 3e**). A clear concentration dependent increase in fluorescence was observed with increasing levels of the acyl acceptor substrate. Finally, we tested various enzyme concentrations and found that there was apparent saturation in the higher enzyme concentrations (**Fig. 3f**). Combined, these results establish the feasibility of conjugating a synthetic, chemically defined acyl donor substrate to a flow cytometry-compatible bead and analyzing crosslinking with a synthetic, chemically defined acyl acceptor substrate.

To investigate the potential for multiplexing applications, we tested the flexibility of this mode of assay with other dyes. The fluorescein dye on the *N*-terminal was replaced with tetramethylrhodamine (**Tm_acceptor**) (**Fig. S2**). Incubation of the TAMRA-labeled acyl acceptor strand with bead-bound **tetAcK-yne** and Ldt_Mt2_ led to a pronounced fluorescence increase over background. As seen with **qSeDAPtri**, inclusion of meropenem completely abolished signal from the **Tm_acceptor** probe, confirming enzyme-dependent fluorescence generation. We next examined the effect of temperature, noting that Ldt_Mt2_ operates at human physiological temperature.^81^ Conducting reactions at 37 °C under optimized conditions produced robust, statistically significant signal comparable to room temperature results (**Fig. S3**), indicating minimal temperature effect on Ldt activity and confirming assay robustness under physiological conditions.

### Expanding detection modalities: Antibody-based readouts

Having established the bead-based fluorescence assay, we next explored the possibility of using an orthogonal detection method to enhance versatility. Because fluorescein is a well-characterized antigen with high-affinity antibodies,^82, 83^ we hypothesized that anti-fluorescein antibody could provide alternative detection of Ldt-mediated crosslinking. Beads were incubated with the **qSeDAPtri** and upon completion of the incubation time with Ldt, beads were treated with anti-FITC IgG, which was subsequently detected using an HRP-fused secondary antibody. Beads were arrayed in standard 96-well plates, and colorimetric signal development was monitored using a microplate reader (**Fig. S4**). The assay produced significant absorbance above background only in the presence of all components. Signal loss upon enzyme omission or meropenem inhibition confirmed feasibility of this ELISA-like format. To eliminate the bead transfer step, we adapted the assay to azide-functionalized microtiter plates. Azide surface density was confirmed by conjugating FAM-alkyne *via* CuAAC, followed by indirect ELISA detection using anti-FITC IgG and HRP-conjugated secondary antibody (**Fig. S5**). Having validated plate functionalization, we immobilized **tetAcK-yne** on well surfaces and performed Ldt-mediated transpeptidation reactions (**Fig. S6a**). Detection with anti-FITC IgG followed by HRP-conjugated secondary antibody yielded robust colorimetric signal above background (**Fig. S6b**). These results indicated platform compatibility with multiple detection strategies, fluorescent or enzyme-linked, thereby expanding analytical flexibility and high-throughput screening potential.

### Profiling transpeptidase activity across *M. smegmatis* Ldt paralogs

Mycobacterial Ldts are classified into six classes based on conserved sequences and structural motifs, with additional biochemical variations among paralogs.^14, 84^ *Mycobacterium smegmatis* (*M. smegmatis*), a common *Mtb* surrogate, encodes six Ldt paralogs (LdtA–F) compared to five in *Mtb* (Ldt_Mt1–5_), all of which are proposed to contribute to PG integrity.^14, 15, 84, 85^ Previously, isolated muropeptides were used to assess the transpeptidase activity of all 5 *Mtb* Ldts in solution showing 3→3 crosslinking activity for all but one of the paralogs, namely the class 3 Ldt, Ldt_Mt3_.^29^ Additionally, genetic deletions of *M. smegmatis* Ldt paralogs reveal their ability to incorporate fluorescent D-amino acids within the PG or to determine the predominant Ldt within *M. smegmatis*.^14, 85^ Beyond genetic manipulations of Ldt paralogs in *M. smegmatis*, LC/MS has been used to demonstrate differential acylation of all six paralogs by various β-lactams.^84^ As such, we leveraged our optimized bead-based platform to systematically profile the transpeptidase activity of these paralogs *in vitro*. All six *M. smegmatis* Ldts were purified and assessed under optimized conditions. LdtA, LdtB, LdtE, and LdtF generated significant fluorescence signals above background (**Fig. 4a, b, e, f**). Upon pre-treatment of the enzymes with meropenem, bead fluorescence returned to background levels. These results confirmed the Ldt-catalyzed activity of these enzymes for the substrates presented here. Notably, this represents, to our knowledge, the first demonstration of catalytic activity for a class 6 Ldt (LdtF) using purified peptidyl substrates, establishing our platform’s utility for characterizing previously uncharacterized enzymes.

**Figure 4.**
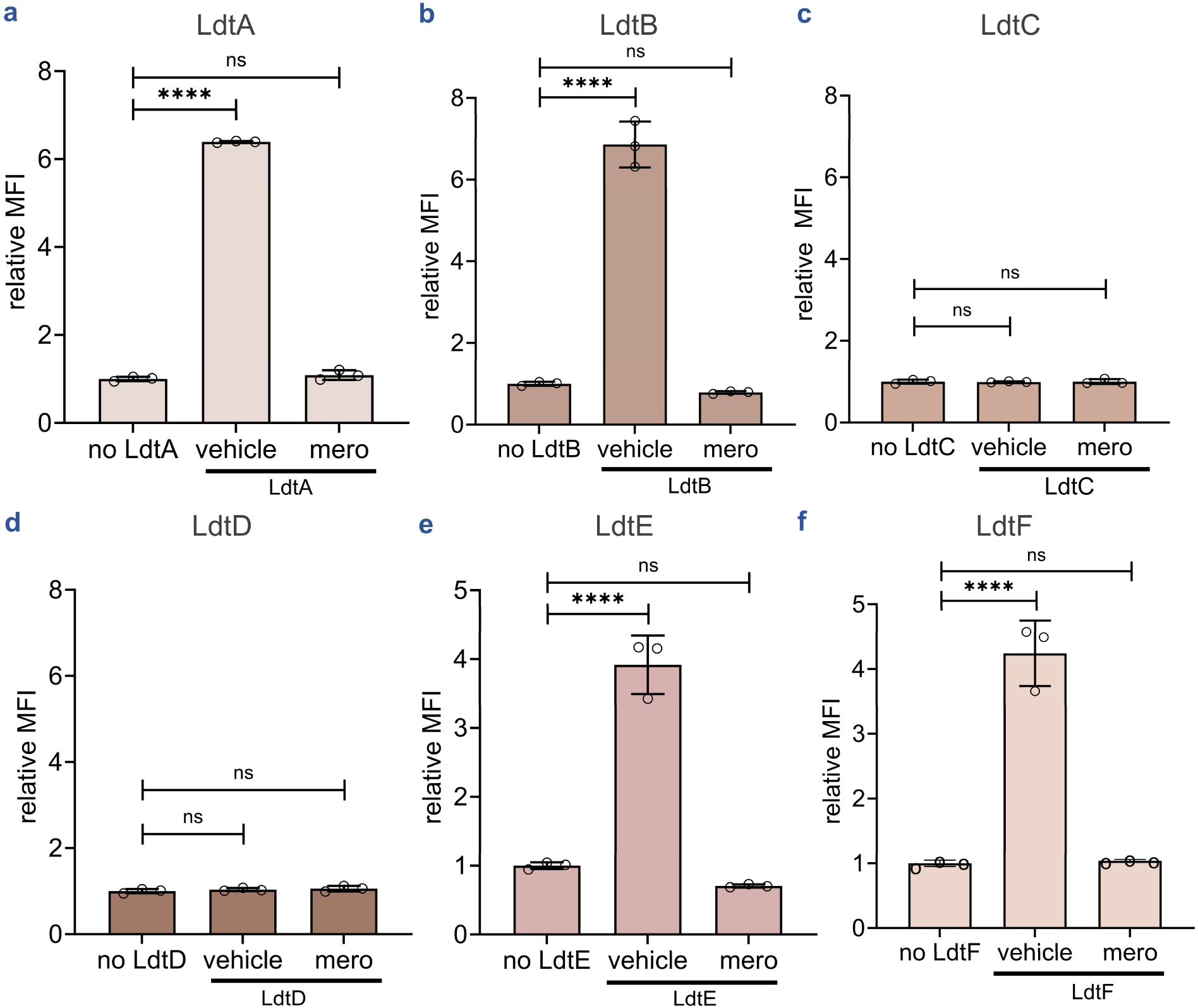
Assessment of transpeptidase activity of *M. smegmatis* Ldt paralogs. Flow cytometry analysis of beads bearing **tetAcK-yne** incubated with **qSeDAPtri** in the presence or absence of LdtA-F for 1h (**a**-**f**). Mean fluorescence intensity (MFI) represents the ratio of fluorescence relative to the control (no Ldt), quantified from 10000 events. Statistical significance was determined by one-way ANOVA (ns = not significant, **** p<0.0001).

Interestingly, no significant signal above background was detected for class 3 enzyme LdtD or class 5 enzyme LdtC (**Fig. 4c, d**). While class 5 Ldt in *Mtb* (Ldt_Mt5_) has been previously reported to form 3→3 crosslinking *in vitro*, and LdtC can function as the sole 3→3 crosslinking enzyme in *M. smegmatis,* to our knowledge there are no reports describing 3→3 crosslinking activity of class 3 Ldts.^15, 29^ Beyond 3→3 PG crosslinking, other Ldts have been found in different species of bacteria to have other roles including anchoring and cleaving of membrane proteins to the PG,^86–91^ unipolar cellular elongation, ^89, 92^ or unusual 1→3 PG crosslink.^93, 94^ In *E. coli*, LdtA (ErfK), LdtB (YbiS), and LdtC (YcfS) primarily catalyzes the covalent attachment of Braun lipoprotein (Lpp) to PG, reinforcing the outer membrane-cell wall linkage.^86^ The class 3 Ldt enzyme (*M.smegmatis* LdtD) utilized here may have a function other than the canonical 3→3 crosslinking being monitored *via* the platform described here. Moreover, LdtD was shown to be acylated by (carba)penems, further hinting at an undefined role that may be inhibited by these antibiotics.^84^ Conversely, the lack of detectable activity by the *M. smegmatis* Class 5 LdtC suggests it cannot efficiently process **qSeDAPtri**, even though its *Mtb* homologue, LdtMt5, forms 3→3 crosslinks *in vitro*.^29^ We hypothesize that this reflects stringent substrate specificity, potentially requiring authentic *m*-DAP-containing acceptors rather than *m*-SeLAN surrogates. Future optimization incorporating *m*-DAP-based probes may elucidate substrate preferences for LdtC (and potentially LdtD) and expand the applicability of the platform for this Ldt class.

### Acyl acceptor profiling reveals chemical modifications tolerated by Ldts

Having established the bead-based assay, we next assessed the breadth of peptide substrates tolerated as acyl acceptors. We have previously shown that tripeptides bearing various residues at the third position function exclusively as acyl acceptors in live bacteria, as the absence of the D-alanyl residue prevents recognition by Ldts to function as a donor stem.^57, 58^ In contrast, tetrapeptides containing a lysine residue at the third position are well tolerated by Ldts and are efficiently incorporated within the PG of live bacteria as either acyl donors or acceptors.^63, 95–97^ Accordingly, our platform, employing a single acyl donor (**tetAcK-yne, Fig. 3a**) and a single acyl acceptor, is well-suited to interrogate how chemical modifications influence Ldt recognition and processing of fluorescent acyl acceptors. While our prior cellular studies provided critical validation of probe uptake and incorporation, these live cell analyses inherently expose PG analogs to the full suite of active transpeptidases and a vast diversity of acyl-complementary ends (**Fig. 2e**). This environmental complexity makes it nearly impossible to systematically dissect how specific structural motifs drive enzymatic function.

To rigorously map the determinants of substrate recognition in a controlled environment, we synthesized a library of eight additional stem peptide analogs that reflect the natural heterogeneity of mycobacterial PG (**Fig. 5a**). These probes were designed to systematically evaluate modifications critical to cell wall architecture, including the amidation status of the *iso*-D-glutamic acid and the identity of the third-position diamino acid (contrasting *m*-DAP with lysine, a hallmark distinction between mycobacterial and other bacterial chemotypes). We also investigated the impact of peptide chain length on acyl-acceptor competence by testing trimeric, tetrameric, and pentameric variants. As before, we initiated our evaluation with Ldt_Mt2_ to establish a baseline for catalytic tolerance. Evaluation of trimeric substrate revealed that Ldt_Mt2_ tolerated only those harboring an *m*-DAP mimetic at the third position, **qSeDAPtri** and **qCYTtri,** consistent with prior observations in live bacteria.^57, 58^ Importantly, the lack of fluorescent signal from **qSeLYStri** underscores the requirement of a side-chain carboxylate in trimeric substrates to enable a productive engagement within the active site of this enzyme as we previously demonstrated in live mycobacteria.^58^

**Figure 5.**
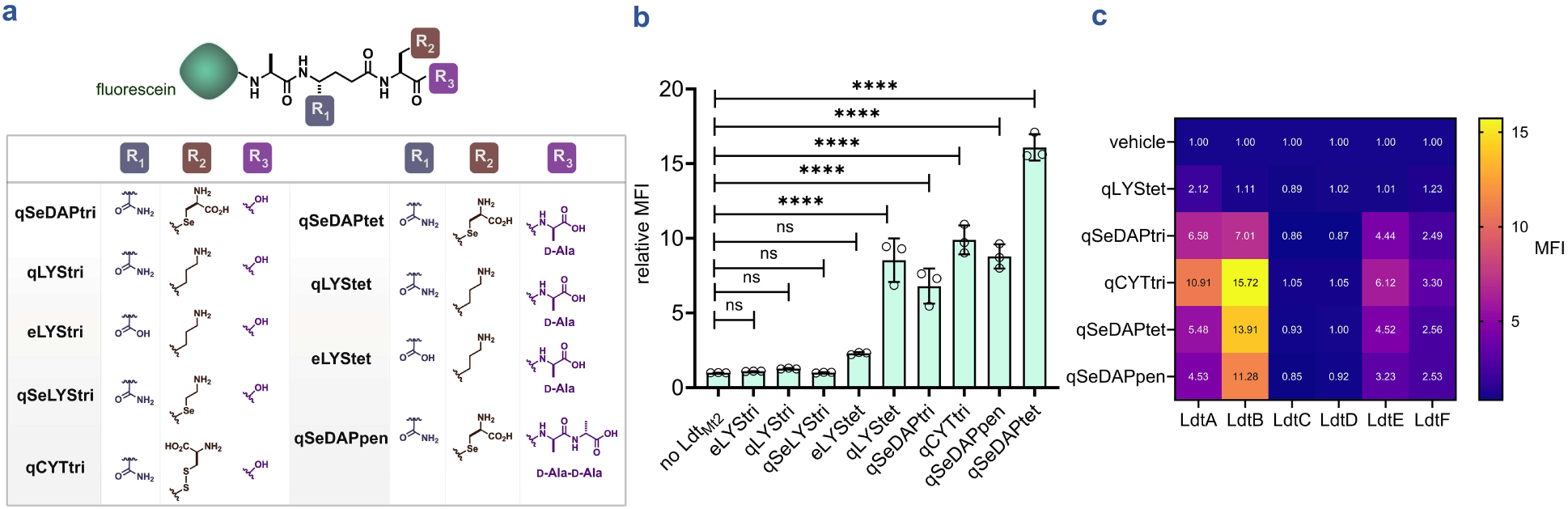
Systematic evaluation of acyl-acceptor tolerance across Ldt enzymes. (**a**) Schematic of acyl acceptor stem peptide analogs used to probe substrate tolerance in *Mtb* Ldt_Mt2_ and across *M. smegmatis* Ldt paralogs. (**b**) Assessment of structural features within acyl acceptors tolerated by Ldt_Mt2_. Fluorescent PG stem peptide analogs with defined compositional differences were evaluated for processing using bead-bearing **tetAcK-yne** and Ldt_Mt2_. Mean fluorescence intensity (MFI) represents the ratio of fluorescence relative to the control (no Ldt), quantified from 10000 events. Statistical significance was determined by one-way ANOVA (ns = not significant, **** p<0.0001). (**c**) Heatmap summarizing the processing of acyl acceptors by *M. smegmatis* Ldt paralogs, as determined using the bead-based assay under optimized conditions. Data represent the mean values from three independent replicates (n=3). P-values were determined by one-way ANOVA (ns = not significant, **** p<0.0001).

We have previously established that fluorescent and bioorthogonal tetrapeptides bearing a third-position lysine are incorporated into the PG of mycobacteria.^63, 95, 96^ Consistent with these findings, assessment of tetrameric acyl acceptors in our assay revealed that Ldt_Mt2_ tolerated all tetrameric substrates except **eLYStet**, which we attribute to the lack of amidation of the *iso*-D-glutamic acid at the second position. Together, these results underscore the importance of *iso*-D-glutamic acid amidation in Ldt processing and highlight the critical role of the terminal D-alanine in substrate recognition and enzymatic catalysis (**Fig. 5b**).^45–47, 63^ Finally, we observed that **qSeDAPpen** was also tolerated by Ldt_Mt2_. We hypothesize that this tolerance arises from the presence of the *m*-DAP mimetic at the third position, despite the substrate being terminated by D,D-stereocenters that preclude Ldt engagement and its function as a donor stem (**Fig. 5b**).^63^

Having established the substrate scope of Ldt_Mt2_, we next evaluated the tolerance of these analogs across *M. smegmatis* Ldt paralogs. Peptides bearing a third-position *m*-DAP mimetic were well tolerated by LdtA, LdtB, LdtE, and LdtF (**Fig. 5c, S7**). Only LdtA tolerated the tetrameric substrate, harboring a third-position lysine (**Fig. 5c, S7).** In contrast, LdtC and LdtD failed to process any of the tested substrates. These results suggest more stringent substrate requirements for crosslinking and support their proposed roles in processes distinct from canonical PG peptide crosslinking. Collectively, these results demonstrate the utility of our platform to systematically interrogate structural modification tolerance across Ldt paralogs.

### High-throughput β-lactam profiling validates screening potential

Having shown that our assay reports on Ldt-mediated crosslinking across multiple enzyme paralogs, we applied it to profile β-lactam antibiotic inhibition. To assess suitability for high-throughput screening, we first evaluated assay quality in 96-well format. **TetAcK-yne** beads were incubated with **qSeDAPtri** in the presence or absence of Ldt_Mt2_ to establish positive and negative controls. We calculated a Z’ score of 0.697 (**Fig. 6a, b**), well above the 0.5 threshold indicating adequate signal separation and reliability for screening applications. We then evaluated a representative β-lactam subset including (carba)penems, ampicillin, monobactams, and cephalosporins (**Fig. 6c** and **Fig. S8**). Each enzyme was preincubated with increasing antibiotic concentrations before assessing transpeptidase activity. For *M. smegmatis* Ldts and Ldt_Mt2_, we observed dose-dependent inhibition by meropenem, faropenem, doripenem, biapenem, and tebipenem, whereas ceftriaxone, aztreonam, and ampicillin exhibited minimal inhibition at tested concentrations (≤20 µM) (**Fig. 6d**, **Fig. 6e**, and **S8**). Half-maximal inhibitory concentrations (IC₅₀) for each enzyme-antibiotic pair were calculated and summarized (**Table S1**). Visualization of pIC₅₀ values (negative log IC₅₀) revealed distinct inhibition profiles across the β-lactam classes tested (**Fig. 6f**). These findings align with previous reports that Ldts belonging to class 1, 2, and 4 are preferentially inhibited by the carbapenem tebipenem and the penem faropenem, and that meropenem is the most potent LdtF (class 6) inhibitor, validating the accuracy of the platform.^17, 31, 33, 34, 84^ Finally, further demonstrating our platform’s reliability is the observed minimal activity of the penam ampicillin, the monobactam aztreonam, and the cephalosporin ceftriaxone within the tested concentration range as these are known to be poor Ldt inhibitors.^33, 34^

**Figure 6.**
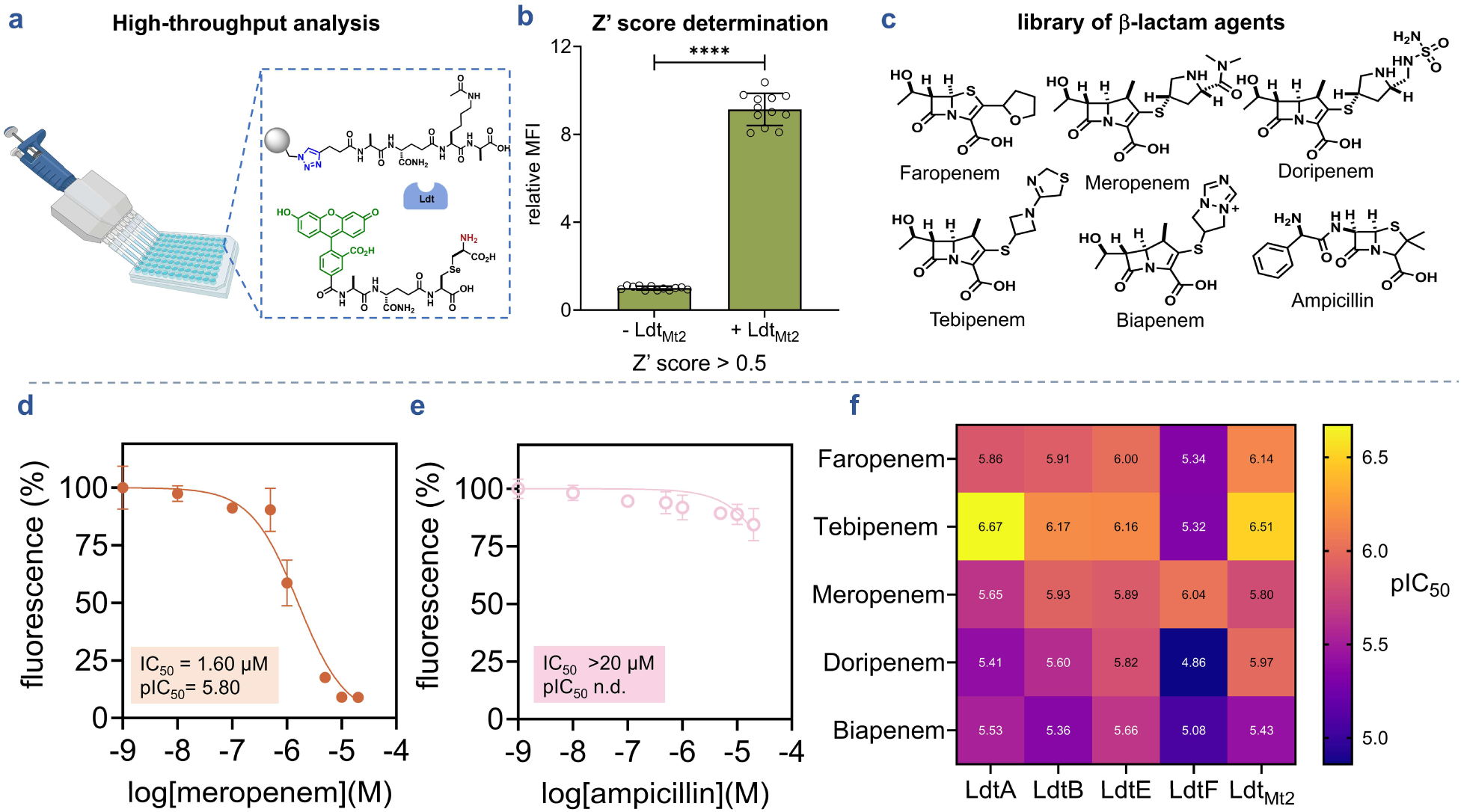
High-throughput bead-based profiling of antibiotic activity against L,D-transpeptidases. (**a**) Application of the bead-based assay platform for high throughput screening. (**b**) Assessment of the separation between the positive (+ Ldt_Mt2_) and the negative control (- Ldt_Mt2_) for Z’ score determination. The calculated Z’ score was 0.697 above the 0.5 threshold, indicating reliable signal separation for screening applications (**c**) Chemical structure of antibiotics spanning multiple β-lactams subclasses including (carba)penem, and penam. (**d**) Representative IC_50_ curve illustrating concentration-dependent inhibition of Ldt_Mt2_ by meropenem using the bead-based assay. (**e**) Lack of inhibitory activity of ampicillin against Ldt_Mt2_ under the same assay conditions. (**f**) Heatmap of pIC_50_ values summarizing inhibition of *M. smegmatis* Ldt paralogs and *Mtb* Ldt_Mt2_ by (carba)penems, determined using the bead-based assay under optimized conditions. Data points represent the mean values of three replicates (n=3).

## CONCLUSION

We have developed a versatile, high-throughput platform for investigating L,D-transpeptidase enzyme activity and inhibition. The system employs a bead-based format where an acyl donor peptide substrate is immobilized on polystyrene beads, and enzyme-catalyzed incorporation of a fluorescent acyl acceptor tripeptide is quantitatively monitored by flow cytometry. This approach is readily adaptable to alternative detection modalities, including indirect ELISA configurations utilizing antibody-based readouts or immobilization of acyl donor at microtiter plate surfaces with antibody-mediated chemiluminescent detection. This flexibility enhances throughput and scalability for diverse screening applications.

Using this platform, we systematically profiled β-lactam antibiotic inhibition across multiple Ldt paralogs, including the primary transpeptidase of *Mtb,* Ldt_Mt2_. Results confirmed that (carba)penems exhibit potent, concentration-dependent inhibition, thereby validating platform accuracy. Beyond inhibitor profiling, the system enabled evaluation of the structure–activity relationships of acyl-acceptor peptides across Ldt paralogs, and characterization of transpeptidase activity across *M. smegmatis* Ldt paralogs, revealing, catalytic activity of the class 6 enzyme LdtF with purified substrates. Conversely, lack of detectable activity for LdtC suggests stringent substrate specificity, highlighting opportunities for future probe optimization.

This platform overcomes critical limitations of existing Ldt activity assays, eliminating dependence on laborious sacculi preparation while maintaining mechanistic fidelity to the native two-step transpeptidation reaction. The demonstrated compatibility with multiple detection modalities, robust performance under physiological conditions, and validated screening metrics establish this as a practical tool for accelerating next-generation therapeutic development. As antimicrobial resistance continues to escalate, particularly mycobacterial infections, such tools are essential for discovering novel scaffolds with therapeutic potential against *Mtb* and related pathogens.

## STATISTICAL ANALYSIS

Unless otherwise specified, statistical analysis was conducted using GraphPad Prism 9.5. Experiments were conducted in triplicate and significant experiments were repeated at least twice. One-way ANOVA was used to calculate statistical significance. Error bars represent standard deviation.

## SUPPORTING INFORMATION

The Supporting Information contains additional figures illustrating experimental data, complete synthetic routes for peptide and small molecule preparation, full characterization data (HRMS, HPLC) for all synthesized compounds, and supplementary references cited in the methodology and discussion.

## ACKNOWLEDGEMENT

This study was supported by the NIH grant 1R01AI178975-01 (M.M.P.), R35GM124893 (M.M.P.), R01AI179080-01 (M.M.P.).

