## Supplementary Information for "Activity Profiling of Mycobacterial L,D-Transpeptidases"

### TABLE OF CONTENTS


| | |
|---|---|
| **Supporting Figures & Table** | **S4-S12** |
| **Figure S1:** Flow cytometry analysis of beads conjugated to Fam-alkyne *via* CuAAC | S4 |
| **Figure S2:** Flow cytometry analysis of $Ldt_{Mt2}$-catalyzed crosslinking of qSeDAPtri onto tetAcK-yne beads | S5 |
| **Figure S3:** $Ldt_{Mt2}$ crosslinking at physiological temperature | S6 |
| **Figure S4:** Antibody-based detection of fluorescein on qSeDAPtri crosslinked beads | S7 |
| **Figure S5:** Antibody-based colorimetric detection of fluorescein on Fam-alkyne well plates | S8 |
| **Figure S6:** Assessment of Ldt-mediated crosslink formation in an azide-functionalized 96-well plate | S9 |
| **Figure S7:** Systematic evaluation of acyl-acceptor tolerance across *M. smegmatis* | S10 |
| **Figure S8:** Inhibition studies of *M. smegmatis* Ldt paralogs and $Ldt_{Mt2}$ | S11 |
| **Table S1:** Calculated $IC_{50}$ and $pIC_{50}$ values | S12 |
| **Experimental Procedures** | **S13-S21** |
| Materials table | S13-S14 |
| Flow cytometry analysis of isolated peptidoglycan/ sacculi or spherotech polystyrene beads | S15 |
| Peptidoglycan/ sacculi isolation of *M. smegmatis* | S15 |
| Expression & purification of $Ldt_{Mt2}$ | S15 |
| Expression &purification of *M. smegmatis* Ldt paralogs | S15 |
| $Ldt_{Mt2}$ catalyzed crosslinking of qSeDAPtri on *M. smegmatis* sacculi | S15-S16 |
| Copper-catalyzed Azide Alkyne Cycloaddition (CuAAC) of Fam-alkyne onto SPHERO azido Polystyrene particles | S16 |
| Copper-catalyzed Azide Alkyne Cycloaddition (CuAAC) of tetAcK-yne onto SPHERO azido Polystyrene particles | S16 |
| Determination of the optimal time needed for crosslinking on beads | S116 |
| Determination of the optimal concentration of tetAcK-yne | S16-S17 |
| Determination of the optimal concentration of qSeDAPtri | S17 |
| Determination of optimal enzyme concentration | S17 |

| | |
|---|---|
| Ldt-catalyzed crosslinking of tetAcK-yne with qSeDAPtri | S18 |
| Assessment of transpeptidation at physiological temperature | S18 |
| Plate reader-based detection of crosslink formation on the bead | S19 |
| Plate reader-based detection of fluorescein conjugated at the bottom of well plates | S19 |
| $Ldt_{Mt2}$ – catalyzed crosslinking at the bottom of well plate | S20 |
| Bead-based crosslinking of peptide substrates by *M. smegmatis* Ldts | S20 |
| Bead-based crosslinking of acyl acceptor analogues with *Mtb* $Ldt_{Mt2}$ and *M. smegmatis* Ldts | S20 |
| Z' score determination | S21 |
| Antibiotics inhibition of *M. smegmatis* Ldts and *Mtb* $Ldt_{Mt2}$ | S21 |
| **Synthesis and Characterization** | **S22-S38** |
| General peptide synthesis and characterization | S22 |
| **References** | **S39** |

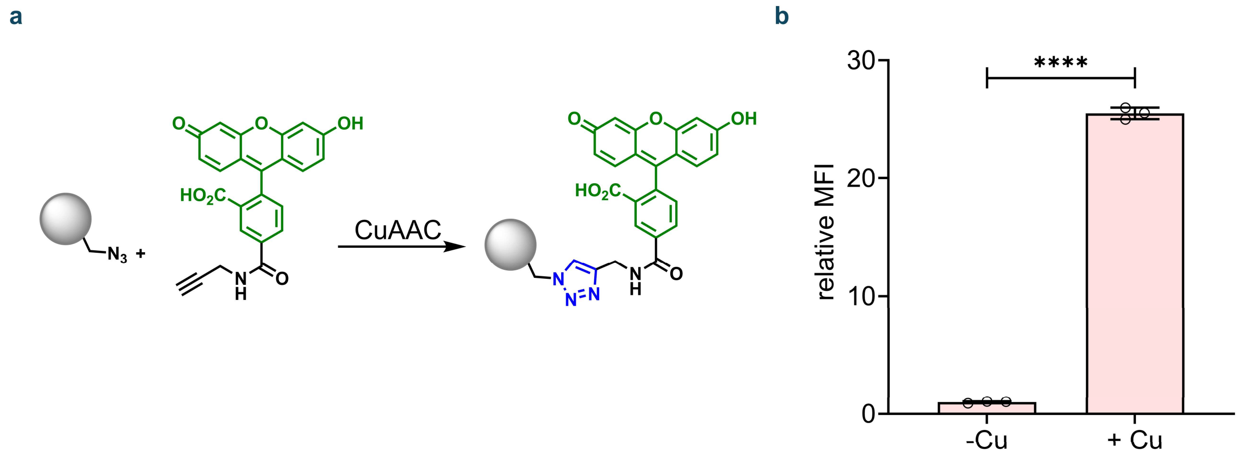


**Figure S1.** Flow cytometry analysis of beads conjugated via copper-catalyzed azide alkyne cycloaddition (CuAAC) with 5 μM fluorescein alkyne (**Fam-alkyne**) for 1h at room temperature, then washed (3X) with PBS containing 0.01% Tween-20. Mean fluorescence intensity (MFI) is the ratio of fluorescence levels above the control (-Cu) treatment from 10000 events. p-values were determined by a two-tailed t-test (* denotes a p-value < 0.05, ** < 0.01, ***<0.001, ns = not significant).

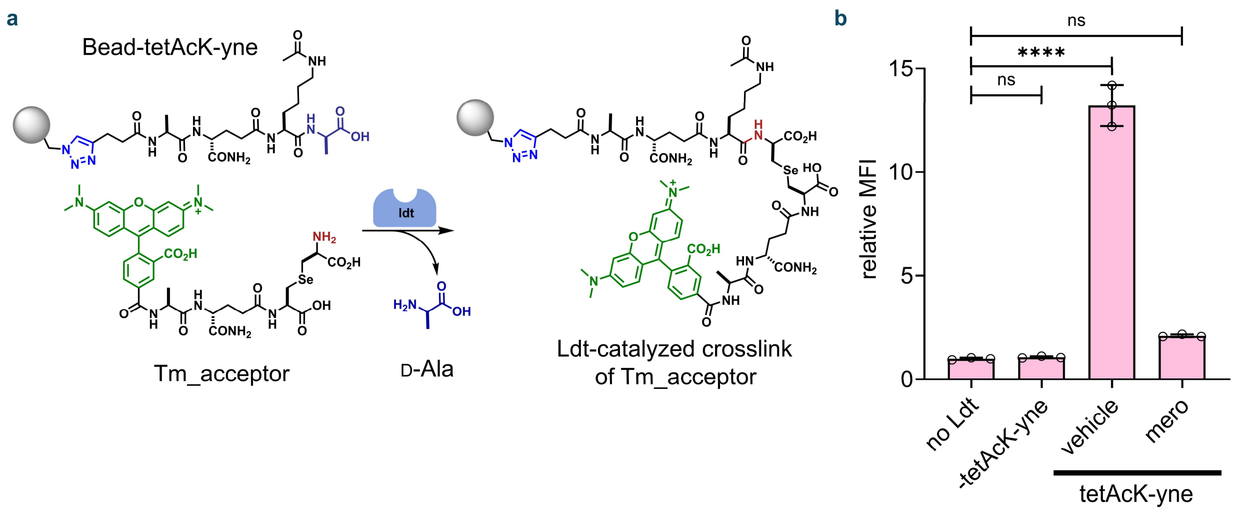


**Figure S2.** Flow cytometry analysis of beads bearing 100 µM of **tetAcK-yne** incubated with 10 µM **qSeDAPtri** in the presence, absence, or inhibition of 1 µM of $Ldt_{Mt2}$ for 1h at room temperature, then washed (3X) with PBS containing 0.01% Tween-20. Mean fluorescence intensity (MFI) is the ratio of fluorescence levels above the control ( no $Ldt_{Mt2}$) treatment from 10000 events. p-values were determined by one-way ANOVA (ns = not significant, **** p<0.0001).

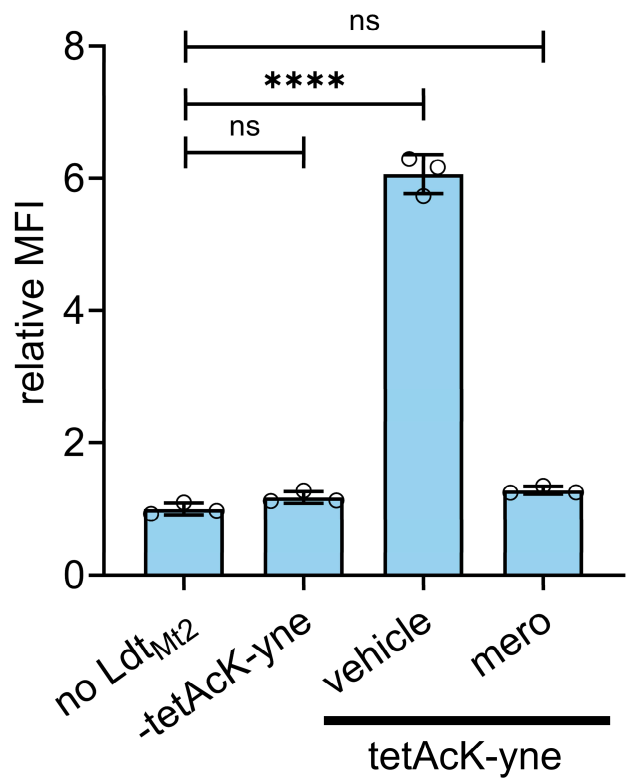


**Figure S3.** Flow cytometry analysis of beads bearing 100 μM of **tetAcK-yne were** incubated with 10 μM **qSeDAPtri,** in the presence or absence of 1 μM of $Ldt_{Mt2}$ for 1h at 37 °C, post which the reaction was quenched with 0.1% TFA and washed (3X) with PBS containing 0.01% Tween-20. Mean fluorescence intensity (MFI) is the ratio of fluorescence levels above the control (no $Ldt_{Mt2}$) treatment from 10000 events. P-values were determined by one-way ANOVA (ns = not significant, **** $p<0.0001$).

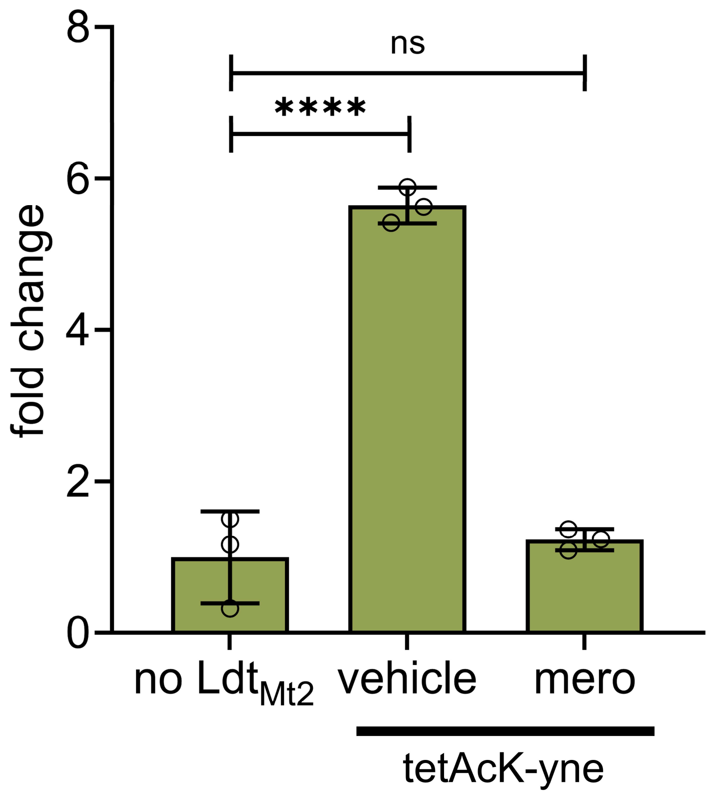


**Figure S4.** Plate reader analysis of **tetAcK-yne** beads crosslinked with **qSeDAPtri** by Ldt$_{Mt2}$. Crosslink formation was assessed using a monoclonal mouse anti-Fluorescein IgG antibody, which binds to the fluorescein moiety on the N-terminus of **qSeDAPtri**. Treatment with an HRP-bearing anti-mouse IgG and treatment with TMB substrate for 30 min and 1M sulfuric acid ($H_2SO_4$) resulted in a signal that was measured at 450nm and 570nm wavelength. Normalized absorbance is the ratio of signal levels above the control beads (no Ldt$_{Mt2}$). P-values were determined by one-way ANOVA (ns = not significant, **** $p<0.0001$).

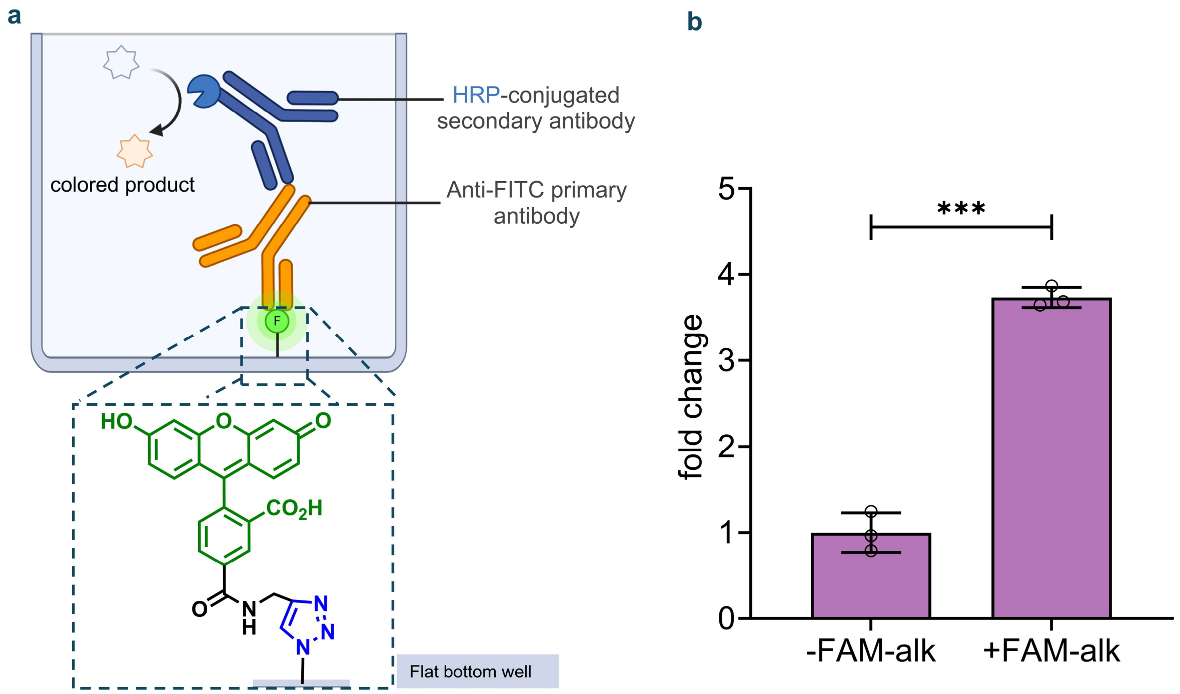


**Figure S5.** Plate reader analysis of 5 μM **Fam-alkyne** conjugated via CuAAC at the bottom of 3D-azido plates. The fluorescein moiety was detected using a monoclonal mouse anti-Fluorescein IgG antibody. Treatment with an HRP-conjugated anti-mouse IgG and treatment with TMB (3,3′,5,5′-tetramethylbenzidine) substrate and 1M sulfuric acid ($H_2SO_4$) resulted in a signal that was measured at both 450nm and 570nm wavelength. Removal of copper from the CuAAC reaction abrogated the colorimetric signal observed. The background signal measured at 570nm was subtracted from the signal measured at 450nm. Normalized absorbance is the ratio of signal levels above the control (-Cu). P-values were determined by a two-tailed t-test (* denotes a p-value < 0.05, ** < 0.01, ***<0.001, ns = not significant.

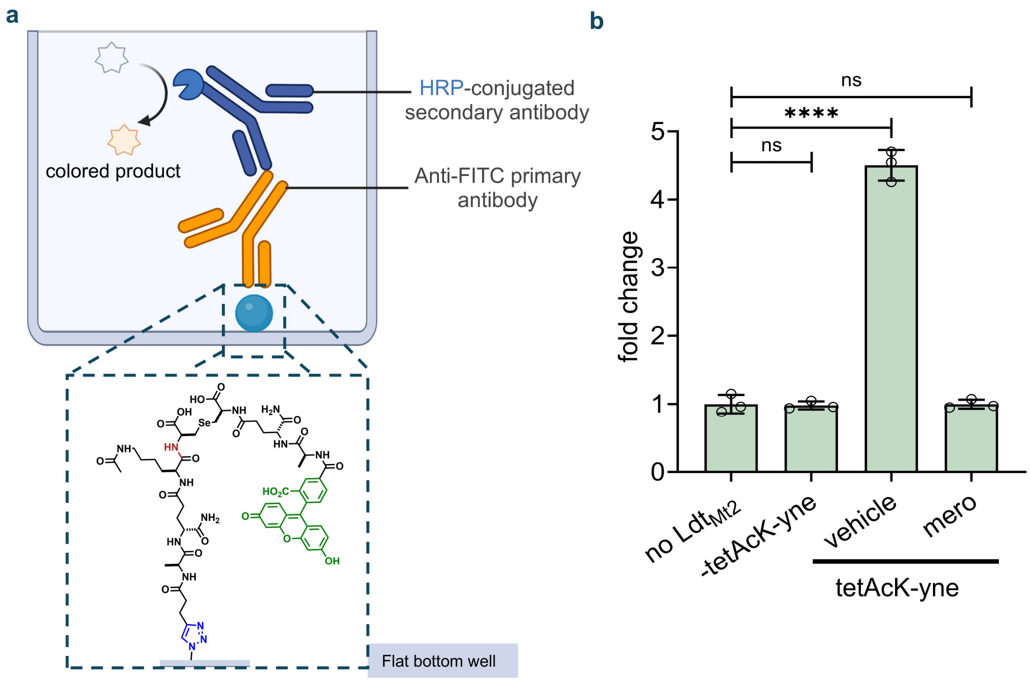


**Figure S6.** (**a**) Schematic representation of the detection of Ldt-mediated crosslink formation in an azide-functionalized 96-well plate. **tetAcK-yne** was immobilized on the bottom of the well via CuAAC, followed by incubation with **QSeDAPtri** and $Ldt_{Mt2}$ to generate a fluorescent crosslink on the surface. The fluorescein moiety was detected using a primary anti-fluorescein antibody, followed by an HRP-conjugated secondary antibody recognizing the Fc region of the primary antibody. Upon addition of TMB (3,3′,5,5′-tetramethylbenzidine), HRP-catalyzed oxidation produced a blue-colored product, which was subsequently quenched with 1M sulfuric acid, and quantified using a plate reader. (**b**) Plate-reader assessment of $Ldt_{Mt2}$-mediated crosslink formation under optimized conditions showed a robust absorbance signal at 450 nm. Omission of $Ldt_{Mt2}$ or its inhibition by meropenem reduced the signal to background levels. Absorbance values are plotted as the ratio of the measured signal relative to the no $Ldt_{Mt2}$ control**.** P-values were determined by one-way ANOVA (ns = not significant, **** p<0.0001).

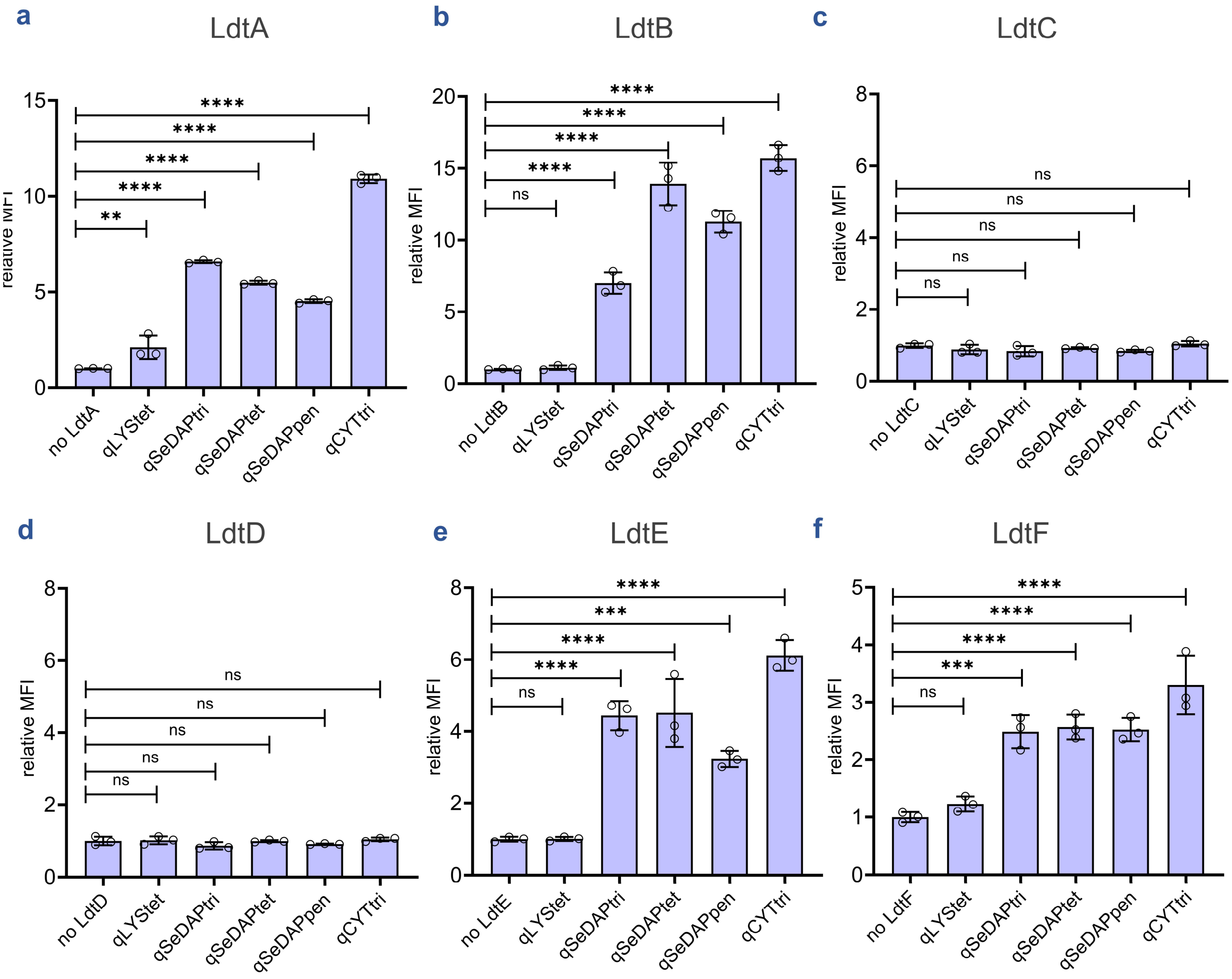


**Figure S7.** Systematic evaluation of acyl-acceptor tolerance across *M. smegmatis*. Flow cytometry analysis of beads bearing **tetAcK-yne** incubated with various acyl acceptor analogues in the presence or absence of LdtA-F for 1h (**a-f**). Mean fluorescence intensity (MFI) represents the ratio of fluorescence relative to the control (no Ldt), quantified from 10000 events. Statistical significance was determined by one-way ANOVA (ns = not significant, **** p<0.0001).

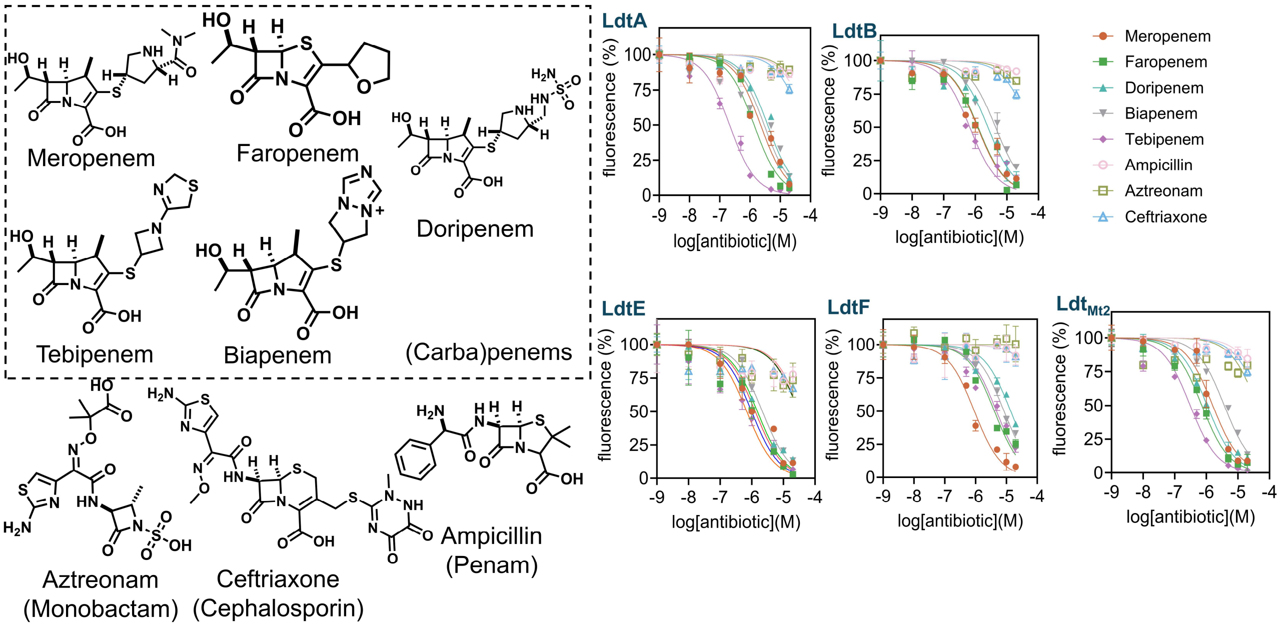


**Figure S8.** Dose response curves of *M. smegmatis* Ldts (A, B, E, F) and *Mtb* $Ldt_{Mt2}$ using the bead-based assay. These assays were run using the bead-based assay under optimized conditions and using a focused library of antibiotics, including penem, carbapenem, monobactam, penicillin, and cephalosporin. The data points displayed represent the mean, with error bars representing the standard error (n=3).

**Table S1. Calculated $IC_{50}$ values**

| | LdtA | LdtB | LdtE | LdtF | $Ldt_{Mt2}$ |
|---|---|---|---|---|---|
| Meropenem | 2.260 µM | 1.186 µM | 1.293 µM | 0.9162 µM | 1.604 µM |
| Faropenem | 1.376 µM | 1.236 µM | 0.996 µM | 4.091 µM | 0.725 µM |
| Doripenem | 3.865 µM | 2.526 µM | 1.515 µM | 1.385 µM | 1.069 µM |
| Biapenem | 2.967 µM | 4.415 µM | 2.185 µM | 8.412 µM | 3.723 µM |
| Tebipenem | 0.213 µM | 0.671 µM | 0.6989 µM | 4.789 µM | 0.310 µM |
| Ampicillin | >20 µM | >20 µM | >20 µM | >20 µM | >20 µM |
| Aztreonam | >20 µM | >20 µM | >20 µM | >20 µM | >20 µM |
| Ceftriaxone | >20 µM | >20 µM | >20 µM | >20 µM | >20 µM |
| **$pIC_{50}$ values calculated using the negative log of the $IC_{50}$** | | | | | |
| Meropenem | 5.646 | 5.926 | 5.889 | 6.038 | 5.795 |
| Faropenem | 5.862 | 5.908 | 6.001 | 5.388 | 6.139 |
| Doripenem | 5.413 | 5.598 | 5.820 | 4.859 | 5.971 |
| Biapenem | 5.528 | 5.355 | 5.660 | 5.075 | 5.429 |
| Tebipenem | 6.672 | 6.173 | 6.156 | 5.320 | 6.508 |

#### Experimental Procedures

##### Materials

| Reagents | Vendors | Catalog number |
|---|---|---|
| **2- Chlorotrityl Chloride resin (1.0-1.6 meq/g, 100-200mesh)** | Chem Impex | 03498 |
| **α-N-Fmoc-amino acids** | Chem Impex | Variables |
| **5,6-Carboxyfluorescein,<br><br>5,(6)-Carboxy-tetramethylrhodamine** | Chem Impex | 00472, 14719 |
| **N,N'-Diisopropylcarbodiimide (DIC)** | Chem Impex | 00110 |
| **Ethyl Cyano(hydroxyamino)acetate (oxymaPure)** | TCI Chemicals | E0847 |
| **HBTU** | Millipore Sigma | 12804 |
| **N,N'-Diisopropylethylamine** | Millipore Sigma | 387649 |
| **Dichloromethane Anhydrous and ACS-grade** | Millipore Sigma | 270997, D65100 |
| **N,N-Dimethylformamide ACS-grade** | Thermofisher | 039117.M6 |
| **Piperidine** | Millipore Sigma | 104094 |
| **Trifluoroacetic acid reagent grade** | AK Scientific | J51721 |
| **Triisopropylsilane** | Chem Impex | 01966 |
| **Methanol ACS-grade** | Millipore Sigma | 179337 |
| **Diethyl ether** | Millipore Sigma | 673811 |
| **2,2-dithiobis(5-nitropyridine) DTNP** | Millipore sigma | 158194 |
| **Thioanisole** | Millipore Sigma | T28002 |
| **Dithiothreitol (DTT)** | Millipore Sigma | D0632 |

| | | |
|---|---|---|
| **Acetonitrile HPLC grade** | Millipore Sigma | 34851 |
| **Trifluoroacetic acid HPLC grade** | Millipore Sigma | 302031 |
| **Meropenem trihydrate** | TCI chemicals | M2279 |
| **Doripenem hydrate** | Cayman Chemical | 16934 |
| **Tebipenem hydrate** | Cayman Chemical | 33875 |
| **Biapenem** | Cayman Chemical | 28822 |
| **Aztreonam** | Cayman Chemical | 19784 |
| **Ceftriaxone sodium salt hydrate** | Cayman Chemical | 18866 |
| **Faropenem** | Millipore Sigma | F8182 |
| **Ampicillin Sodium salt** | Millipore Sigma | A9518 |
| **Azide click chemistry polystyrene particles** | Spherotech | ACCP-50-5 |
| **FAM (fluorescein)-alkyne, 5-isomer** | Lumiprobe | B41B0 |
| **3D-Azide surface chemistry** | PolyAn | 006 95 601 |
| **Hydrofluoric acid (49%)** | Millipore Sigma | 339261 |
| **LB broth (Miller)** | Millipore Sigma | L3522 |
| **DNAse I** | Millipore Sigma | 10104159001 |
| **RNAse A** | Millipore Sigma | 10109142001 |
| **Trypsin** | Millipore Sigma | T6567 |
| **Sodium dodecyl sulfate (SDS)** | TCI chemicals | I0352 |
| **HRP goat anti-mouse IgG** | Biolegend | 405306 |
| **Monoclonal mouse Anti-Fluorescein (FITC) IgG** | Jackson Immuno Research | 200-002-037 |

#### Flow cytometry analysis of isolated peptidoglycan/ sacculi or spherotech polystyrene beads

Isolated peptidoglycan and spherotech beads from experiments below were analyzed using an Attune NxT flow cytometer equipped with a 488 nm and a 561nm laser with 525/40 nm and 585/16nm bandpass filters. The data were analyzed using the Attune NxT Software, where populations were gated, and no less than 10,000 events per sample were recorded. Wherever presented, the mean fluorescence intensity (MFI) is the ratio of fluorescence levels above the control treatment.

#### Peptidoglycan/ sacculi isolation of *M. smegmatis*

Methodology for isolation of sacculi is an adaptation from published protocols.[1-4] *Mycobacterium smegmatis* strains mc2 155, ATCC 14468 were grown in 7H9 media with 0.5% glycerol, 0.05% tween 80, and 1x ADC enrichment (10 x ADC, 5 g bovine serum albumin, 2 g dextrose, 3 mg catalase in 100 mL deionized water, sterilized by filtration through a 0.2 µm filter before use). *M smegmatis* was inoculated from the glycerol stock by 1 to 1000 dilution to 3 mL fresh 7H9 media with ADC and grown to the stationary phase. Then, cell pellets were resuspended in 200 µL 10 mM NH4HCO3 with protease inhibitor and bath sonicated for 30 min. 10 µg/mL DNase and RNase were added to each well directly after sonication, and the tube was placed in 4 $^{o}$C for 1 h. The cell wall-enriched fractions were collected by centrifugation at 2700 g for 10 min. The pellets were then treated with 200 µL PBS with 2% sodium dodecyl sulfate (SDS) and incubated at 50 $^{o}$C for 1 h with shaking at 250 rpm thrice. Finally, the suspension was centrifuged, and the resulting pellet was reconstituted in 200 µL PBS with 1% SDS and 0.1 mg/mL proteinase K, incubated at 45 $^{o}$C for 1h with shaking at 250 rpm. The tube with the resulting suspension was then heated with boiling water for 1 h and then centrifuged at 2700 g for 10 min. The supernatant was discarded, and the 1%SDS boiling step was repeated 2 times and centrifuged. The pellets were then washed twice with PBS and 4 times with deionized water to give mycolyl-arabinogalactanpeptidoglycan Complex (MAPc). The pellet was resuspended in 200 µL methanol with 0.5%(w/v) KOH and incubated in 37 $^{o}$C at 250 rpm for 4 days. The suspension was then centrifuged, and the pellets were washed with methanol twice and diethyl ether twice and air-dried to afford arabinogalactanpeptidoglycan (AGPG). The resulting AGPG was resuspended in 200 µL deionized water. AGPG was digested with 0.05 N H2SO4 at 37 $^{o}$C for 5 days with shaking at 250 rpm, centrifuged, and washed 4 times with deionized water to give insoluble peptidoglycan (PG). The sacculi were recovered by centrifugation at 21,000 X G for 20 min, washed 5 X with 0.1 M Tris pH 8, resuspended in Milli Q water, and stored at –20 °C for subsequent analysis.

#### Expression of $Ldt_{Mt2}$ proteins and purification

*$Ldt_{Mt2}$* was cloned into a modified pET28a vector, expressed, and purified as we reported previously.[5, 6] The plasmid was transformed into *E. coli* BL21(DE3) cells (NEB). Cultures were grown at 37 °C to an OD600 of ~0.5, cooled to 16 °C, induced with 100 µM IPTG, and incubated with shaking for 20 h. Cells were harvested by centrifugation (3500 × g, 10

min, 4 °C), stored at –20 °C, and then resuspended in lysis buffer (25 mM Tris-HCl, pH 8.0, 400 mM NaCl, 10% glycerol, 1 mM TCEP, protease inhibitor cocktail). After sonication, lysates were clarified by centrifugation (24,500 × g, 30 min, 4 °C) and applied to Ni-NTA resin for 1 h at 4 °C. Bound protein was eluted over a 20–500 mM imidazole gradient, and fractions containing $His_6$-$Ldt_{Mt2}$ were pooled. For tag removal, pooled fractions were dialyzed for 48 h at 4 °C against 25 mM Tris-HCl (pH 8.0), 100 mM NaCl, 10% glycerol, and 1 mM TCEP in the presence of TEV protease (1:100 w/w). Dialysis buffer was exchanged three to four times. The digest was reapplied to Ni-NTA resin, and the flow-through containing tag-free $Ldt_{Mt2}$ was collected. Protein purity was verified by SDS-PAGE, concentrations were determined spectrophotometrically, and aliquots were flash frozen in liquid nitrogen and stored at –80 °C.

##### Expression of *M. smegmatis* Ldts

*M. smegmatis* Ldts were cloned into a modified pET28b vector, expressed, and purified as we reported previously.[7] Briefly, *E. coli* BL21(DE3) cells harboring Ldt-pET28b constructs were cultured in LB medium at 20 °C with shaking until $OD_{600}$ reached ~0.5, at which point protein expression was induced with 100 µM IPTG for 14–20 h. Cells were collected by centrifugation, resuspended in buffer (25 mM Tris, pH 8.0, 400 mM NaCl, 10% glycerol), and lysed by sonication. After clarification, lysates were applied to Ni-NTA resin, and bound proteins were eluted with imidazole. Fractions containing Ldt were identified by SDS-PAGE, pooled, and dialyzed overnight against 50 mM Tris (pH 8.0), 100 mM NaCl, and 10% glycerol in the presence of TEV protease (1:100, w/w). Cleaved proteins were separated from the $His_6$-tag and TEV protease by a second Ni-NTA step, followed by dialysis into storage buffer (50 mM Tris, pH 8.0, 100 mM NaCl, 10% glycerol, 1 mM TCEP). Proteins were concentrated to ≥1 mg/mL, flash frozen in liquid nitrogen, and stored at −80 °C.

##### $Ldt_{Mt2}$ catalyzed crosslinking of qSeDAPtri on *M. smegmatis* sacculi

Sacculi were pipetted into wells of a round-bottom, untreated 96-well plate containing 25 mM Tris pH 8, 2µM $ldt_{Mt2}$, and 20 µM **qSeDAPtri** at room temperature for 5h. Control wells excluded the enzyme. This experiment was performed in triplicate. Following incubation, the enzymatic reaction was quenched using 0.1% trifluoroacetic acid (TFA). The sacculi were then washed thrice by centrifugation at 4000 X g for 3min with PBS containing 0.01% Tween-20. Sacculi fluorescence was assessed by an Attune NxT flow cytometer using the 488nM laser and the 525/49 bandpass filter.

##### Copper-catalyzed Azide Alkyne Cycloaddition (CuAAC) of Fam-alkyne onto SPHERO azido Polystyrene particles

Sphero azido polystyrene beads 3.31 um 5% w/v (250 µL) were resuspended in 1X PBS in a round-bottom untreated 96-well plate then treated with 5 µM **Fam-alkyne**, 1 mM $CuSO_4$, 128 µM THPTA, 1.2 mM L-Ascorbic Acid. The negative control included all the reagents except for $CuSO_4$. The particles were incubated at 37 °C for 2 h. After incubation, they were washed 3X by centrifugation at 4000 X g for 3min with 1X PBS containing 0.01% Tween-20. Bead fluorescence was assessed by an Attune NxT flow cytometer using the 488nM laser and the 525/49 bandpass filter as described previously

**Copper-catalyzed Azide Alkyne Cycloaddition (CuAAC) of tetAcK-yne onto SPHERO azido Polystyrene particles**

Sphero azido polystyrene beads 3.31 um 5% w/v (250 µL) were resuspended in 1X PBS in Eppendorf tubes, then treated with 100 µM purified acyl donor peptide **tetAcK-yne** (see scheme below), 1 mM $CuSO_4$, 128 µM THPTA, 1.2 mM L-Ascorbic Acid. The particles were incubated at 37 °C for 2 h. After incubation, they were washed 3X by centrifugation at 4000 X g for 3min with 1X PBS containing 0.01% Tween-20. They were resuspended to a 5% w/v suspension in 1X PBS.

**Determination of the optimal time needed for crosslinking on beads**

In a round-bottom untreated 96- well plate, 5 µl of beads bearing 100 µM of **tetAcK-yne** were incubated with 25 mM Tris pH 8, 2µM $Ldt_{Mt2}$, and 20 µM **qSeDAPtri** at room temperature for 1,5,10,15,20,25,30,45,60,90, and 120 minutes. The beads were thoroughly mixed by pipetting prior to incubating them with shaking at room temperature for the indicated time. Each condition was run in triplicate. The addition of 0.1% TFA was used to stop enzymatic catalysis at each time point. The beads were subsequently washed thrice by centrifugation at 4000 X g for 3min with PBS containing 0.01% Tween-20. Bead fluorescence was assessed by an Attune NxT flow cytometer using the 488nM laser and the 525/49 bandpass filter as described previously.

**Determination of the optimal concentration of tetAcK-yne**

Sphero azido polystyrene beads 3.31 µm 5% w/v (Spherotech, 50 µL) were washed in PBS and varying concentrations of purified acyl donor peptide **tetAcK-yne** (50,100, 500, and 1000 µM ) was reacted with the azido beads in PBS containing 1 mM $CuSO_4$, 128 µM THPTA, 1.2 mM L-Ascorbic Acid. The particles were incubated at 37 °C for 2 h. After incubation, they were washed thrice with 1X PBS and were resuspended to a 5% w/v suspension. Then, in a round-bottom untreated 96-well plate (VWR cat 82050-622), 5 µl of beads bearing the appropriate concentration of **tetAcK-yne** were incubated with 25 mM Tris pH 8, 2µM $Ldt_{Mt2}$, and 20 µM **qSeDAPtri**. The beads were thoroughly mixed by pipetting prior to incubating them with shaking at room temperature for 1h. Each of the conditions here was run in triplicate. Negative controls were azido beads free of the acyl donor peptide. The addition of 0.1% TFA stopped enzymatic catalysis after the incubation period. The beads were subsequently washed thrice by centrifugation at 4000 X g for 3min with PBS containing 0.01% Tween-20. Bead fluorescence was assessed by an Attune NxT flow cytometer using the 488nM laser and the 525/49 bandpass filter as described previously.

**Determination of the optimal concentration of qSeDAPtri**

In a round-bottom, untreated 96-well plate, 5 µl of beads bearing 100 µM of **tetAcK-yne** were incubated with 25 mM Tris pH 8, 2µM $Ldt_{Mt2}$, and varying concentrations of **qSeDAPtri**, respectively 0.1, 0.5,1,5,10, and 20 µM. The beads were thoroughly mixed by pipetting prior to incubating with shaking at room temperature for 1h. The addition of 0.1% TFA stopped enzymatic catalysis after the incubation period. Each condition tested here was done in triplicate. Negative control wells excluded the addition of the enzyme. The beads were subsequently washed thrice by centrifugation at 4000 X g for 3min with PBS containing 0.01% Tween-20. Bead fluorescence was assessed by an Attune NxT flow cytometer using the 488nM laser and the 525/49 bandpass filter as described previously.

**Determination of optimal enzyme concentration**

In a round-bottom, untreated 96-well plate, 5 µl of beads bearing 100 µM of **tetAcK-yne** were incubated with 25 mM Tris pH 8, 10 µM of **qSeDAPtri**, and varying concentrations of $ldt_{Mt2,}$ respectively 0.1, 0.5,1, and 2 µM of $Ldt_{Mt2}$.The beads were thoroughly mixed by pipetting prior to incubating with shaking at room temperature for 1h. Negative control wells included all but the enzyme. The addition of 0.1% TFA stopped enzymatic catalysis after the incubation period. The beads were subsequently washed thrice by centrifugation at 4000 X g for 3min with PBS containing 0.01% Tween-20. Bead fluorescence was assessed by an Attune NxT flow cytometer using the 488nM laser and the 525/49 bandpass filter as described previously.

**Ldt-catalyzed crosslinking of tetAcK-yne with qSeDAPtri**

In a round-bottom, untreated 96-well plate, 5 µl of beads bearing 100 µM of **tetAcK-yne** were incubated with 25 mM Tris pH 8, 10 µM of **qSeDAPtri,** and 1 µM of $Ldt_{Mt2}$. Negative control wells included all but the enzyme. Inhibition of the enzyme was performed through treatment with 20 µM of freshly prepared meropenem. The beads were thoroughly mixed by pipetting prior to incubating them with shaking at room temperature for 1h. The addition of 0.1% TFA stopped enzymatic catalysis after the incubation period. The beads were subsequently washed thrice by centrifugation at 4000 X g for 3min with PBS containing 0.01% Tween-20. Bead fluorescence was assessed by an Attune NxT flow cytometer using the 561nM laser and the 585/16 bandpass filter as described previously.

**Assessment of transpeptidation at physiological temperature**

In a round bottom untreated 96-well plate, 5 µl of beads bearing 100 µM of **tetAcK-yne** were incubated with 25 mM Tris pH 8, 10 µM of **qSeDAPtri ,** and 1 µM of $Ldt_{Mt2}$. Negative control wells included all reagents but the enzyme. Inhibition of the enzyme was performed through treatment with 20 µM of freshly prepared meropenem. The beads were thoroughly mixed by pipetting prior to incubating them with shaking at 37 °C for 1h. The addition of 0.1% TFA stopped enzymatic catalysis after the incubation period. The beads were subsequently washed thrice by centrifugation at 4000 X g for 3min with PBS containing 0.01% Tween-20 (PBST). Bead fluorescence was assessed by an Attune NxT flow cytometer using the 488nM laser and the 525/49 bandpass filter as described previously.

**Plate reader-based detection of crosslink formation on the bead**

In a round-bottom untreated 96-well plate, 20 µl of beads bearing 100 µM of **tetAcK-yne** were incubated with 25 mM Tris pH 8, 10 µM of **qSeDAPtri,** and 1 µM of $Ldt_{Mt2}$. Negative control wells included all reagents but the enzyme. The beads were thoroughly mixed by pipetting prior to incubating them with shaking at room temperature for 1h. The addition of 0.1% TFA stopped enzymatic catalysis after the incubation period. The beads were subsequently washed thrice by centrifugation at 4000 X g for 3min with PBS containing 0.01% Tween-20. Following the reaction, 5 µl of beads were taken, and fluorescence was assessed by an Attune NxT flow cytometer using the 488nM laser and the 525/49 bandpass filter as described previously. The remainder of the beads were blocked using PBST containing 1% Bovine Serum Albumin (BSA) for 1h. Following the blocking, the beads were washed 4 X by centrifugation at 4000 X g for 3min with PBST. Next, the beads were incubated with monoclonal mouse IgG anti-Fluorescein antibody diluted in PBST (1:1000). Then, the beads were washed 4X by centrifugation at 4000 X g for 3min with PBST and subsequently incubated with anti-mouse IgG conjugated with HRP diluted in PBST (1:5000) for 1h. The beads were washed 4X by centrifugation at 4000 X g for 3min with PBST. Then the beads were incubated with TMB substrates for 30 min. The reaction was stopped by the addition of 1M sulfuric acid ($H_2SO_4$). The absorbance was monitored on an Agilent Gen61.04 plate reader at 450nM and 570nM. The absorbance plotted was the difference of the absorbance at the two wavelengths monitored.

**Plate reader-based detection of fluorescein conjugated at the bottom of well plates**

PolyAn 3D-azido plate was treated with 5 µM fluorescein alkyne **(FAM-alkyne)**, along with 1 mM $CuSO_4$, 128 µM THPTA, 1.2 mM L-Ascorbic Acid in 1X PBS at 37 °C for 2 h. Following the CuAAC, the plate was washed thrice with PBST and then blocked using PBST containing 1% Bovine Serum Albumin (BSA) for 1h. Following the blocking, the wells were washed 4X by centrifugation at 4000 X g for 3min with PBST. Next, the wells were incubated with monoclonal mouse IgG anti-Fluorescein antibody diluted in PBST with 1% BSA (1:1000). Then, they were washed 4 X with PBST and subsequently incubated with anti-mouse IgG conjugated with HRP diluted in PBST with 1%BSA (1:5000) for 1h. The wells were washed 4X with PBST. Finally, the wells were incubated with TMB substrates for 30 min. The reaction was stopped by the addition of 1M sulfuric acid ($H_2SO_4$). Wells in which $CuSO_4$ was omitted and wells treated only with the secondary anti-mouse HRP IgG served as controls for this experiment. The absorbance was monitored on an Agilent Gen61.04 plate reader at 450nM and 570nM. The absorbance plotted was the difference of the absorbance at the two wavelengths monitored.

**$Ldt_{Mt2}$ – catalyzed crosslinking at the bottom of the well plate**

PolyAn 3D-azido plates were treated with 100 µM purified acyl donor peptide **tetAcK-yne**, 1 mM $CuSO_4$, 128 µM THPTA, 1.2 mM L-Ascorbic Acid in 1X PBS at 37 °C for 2 h. Following the CuAAC, the plate was washed thrice with 1 X PBS. For the transpeptidation reaction, the wells were incubated with 25 mM Tris pH 8, 10 µM of **qSeDAPtri**, and 1 µM of $Ldt_{Mt2}$ at room temperature for 1h. Negative control wells included all reagents but the enzyme. Inhibition of the enzyme was performed through treatment with 20 µM of freshly prepared meropenem. The addition of 0.1% TFA stopped enzymatic catalysis after the incubation period. The wells were washed thrice with PBST and then blocked using PBST containing 1% Bovine Serum Albumin (BSA) for 1h. Following the blocking, the wells were washed 4X by centrifugation at 4000 X g for 3min with PBST. Next, the wells were incubated with monoclonal mouse IgG anti-Fluorescein antibody diluted in PBST with 1% BSA (1:1000). Then, they were washed 4 X with PBST and subsequently incubated with anti-mouse IgG conjugated with HRP diluted in PBST with 1%BSA (1:5000) for 1h. The wells were washed 4X with PBST. Finally, the wells were incubated with TMB substrates for 30 min. The reaction was stopped by the addition of 1M sulfuric acid ($H_2SO_4$). The absorbance was monitored on an Agilent Gen61.04 plate reader at 450nM and 570nM. The absorbance plotted was the difference of the absorbance at the two wavelengths monitored.

**Bead-based crosslinking of peptide substrates by *M. smegmatis* ldts**

In a round-bottom, untreated 96-well plate, 5 µl of beads bearing 100 µM of **tetAcK-yne** were incubated with 25 mM Tris pH 8, 10 µM of **qSeDAPtri**, and 1 µM of Ldt A-F. Negative control wells included all reagents but the enzyme. Inhibition of the enzyme was performed through treatment with 20 µM of freshly prepared meropenem. The wells were thoroughly mixed by pipetting prior to incubating them with shaking at room temperature for 1h. The addition of 0.1% TFA stopped enzymatic catalysis after the incubation period. The beads were subsequently washed thrice by centrifugation at 4000 X g for 3min with PBS containing 0.01% Tween-20 (PBST). Bead fluorescence was assessed by an Attune NxT flow cytometer using the 488nM laser and the 525/49 bandpass filter as described previously.

**Bead-based crosslinking of acyl acceptor analogues with *Mtb* $Ldt_{Mt2}$ and *M. smegmatis* ldts**

In a round-bottom untreated 96-well plate, 5 µl of beads bearing 100 µM of **tetAcK-yne** were incubated with 25 mM Tris pH 8, 10 µM of **acyl acceptor**, and 1 µM of $Ldt_{Mt2}$, Ldt A-F. Negative control wells included all reagents but the enzyme. The wells were thoroughly mixed by pipetting prior to incubating them with shaking at room temperature for 1h. The addition of 0.1% TFA stopped enzymatic catalysis after the incubation period. The beads were subsequently washed thrice by centrifugation at 4000 X g for 3min with PBS containing 0.01% Tween-20 (PBST). Bead fluorescence was assessed by an Attune NxT flow cytometer using the 488nM laser and the 525/49 bandpass filter as described previously.

**Z' analysis**

In a round-bottom untreated 96-well plate, 5 µl of beads bearing 100 µM of **tetAcK-yne** were incubated with 25 mM Tris pH 8, 10 µM of **qSeDAPtri**, and 1 µM of $Ldt_{Mt2}$ (positive control n=12). Negative control wells (n=12) included all reagents but the enzyme. The wells were thoroughly mixed by pipetting prior to incubating them with shaking at room temperature for 1h. The addition of 0.1% TFA stopped enzymatic catalysis after the incubation period. The beads were subsequently washed thrice by centrifugation at 4000 X g for 3min with PBS containing 0.01% Tween-20 (PBST). Bead fluorescence was assessed by an Attune NxT flow cytometer using the 488nM laser and the 525/49 bandpass filter as described previously. The Z′ score was calculated from replicate measurements of positive and negative control wells as one minus three times the sum of the standard deviations of the positive and negative controls divided by the absolute difference between their mean signals. ( $Z' = 1 - [3(\sigma_p + \sigma_n) / |\mu_p - \mu_n|]$)

**Antibiotic inhibition of *M. smegmatis* ldts and *Mtb* $Ldt_{Mt2}$**

In round-bottom untreated 96-well plates, purified Ldts ( *M.smegmatis* Ldt A,B,E,F and *Mtb* $ldt_{Mt2}$) at a concentration of 1 µM were incubated with 25 mM Tris pH 8 and varying concentrations (0, 0.01, 0.1,0.5, 1,5,10, and 20 µM) of antibiotic for 20 min. The solutions were thoroughly mixed by pipetting prior to incubation. Then, 5 µl of **tetAcK-yne** beads and 10 µM of **qSeDAPtri** were added and allowed to shake for 1h at room temperature**.** The addition of 0.1% TFA stopped enzymatic catalysis after the incubation period. The beads were subsequently washed thrice by centrifugation at 4000 X g for 3min with PBS containing 0.01% Tween-20 (PBST). Bead fluorescence was assessed by an Attune NxT flow cytometer using the 488nM laser and the 525/49 bandpass filter as described previously. The antibiotics tested included meropenem, faropenem, tebipenem, doripenem, biapenem, ceftriaxone, aztreonam, and ampicillin. Each antibiotic solution was freshly prepared in Milli Q water as a 10mg/ml stock solution immediately prior to each experiment and diluted to the desired concentrations. $IC_{50}$ values were determined using GraphPad. Data points were plotted as the mean with standard deviation as the error bars. The values were normalized against the mean average of no-inhibitor controls and the mean average of no-enzyme controls. The dose–response analysis was performed by using the log(inhibitor) vs. normalized response−variable slope model in GraphPad.

### Synthesis & Characterization

#### General procedure for the solid-phase synthesis of all peptides.

All peptides were prepared by standard Fmoc-based solid-phase peptide synthesis (SPPS) using chlorotrityl chloride resin. Briefly, to a 25 mL peptide synthesis vessel, an appropriate amount of resin was added, followed by the first amino acid (1.1 eq) and N,N-Diisopropylethylamine (DIEA, 4.4 eq) in anhydrous dichloromethane (DCM, 5mL). This was followed by shaking at room temperature for 1h. The resin was then washed with methanol (CH3OH) and dichloromethane (DCM) three times. After the last wash, 3 equiv of the next amino acid was added along with 3 equiv. of ethyl cyanohydroxyiminoacetate (OxymaPure) and 3 equiv. of N, N'-Diisopropylcarbodiimide (DIC). The resin was shaken at room temperature for 2 h, then washed with CH3OH/DCM. The remainder of the amino acids were subsequently coupled in a similar fashion. The fluorophore was coupled overnight using 3 eq HBTU and 6eq DIEA in DMF, followed by a wash. Peptides cleavage from the resin was performed using a TFA/TIPS/H2O cocktail (95:2.5:2.5, v/v/v), shaking at room temperature for 2 h. The solution was filtered and concentrated prior to precipitation by the addition of cold diethyl ether to yield crude peptide. Crude peptides were purified by reverse-phase preparative high-performance liquid chromatography (RP-HPLC) equipped with Waters 1525 with 2489 UV/Visible Detector on a Phenomenex Luna 10 µm C18(2) 100 Å (250 x 21.2 mm) column using gradient elution with water/acetonitrile (H2O/MeCN) containing 0.1% TFA at 10 mL/min. The HPLC fractions of the desired purified compounds were first concentrated under reduced pressure using a rotary evaporator. The final concentrated aqueous solutions were lyophilized to dryness using Labconco Freezone 4.5 L (-84 $^{o}$C) lyophilizer. The peptides were analyzed for purity using Phenomenex Luna 5 µm C18(2) on the same RP-HPLC; gradient elution in H2O/MeCN with 0.1% TFA at 1 mL/min. Peptide identities were confirmed via high-resolution electrospray ionization mass spectrometry (HRMS, ESI/MS). Analyses were obtained on an Agilent 6545B Q-TOF LC/MS equipped with 1260 infinity II LC system with auto sampler. Samples were dissolved in CH3CN and eluted with a CH3CN/H2O solution containing 0.1% formic acid

**Scheme S1. Synthesis of qSeDAPtri precursor qSe(mob)tri**

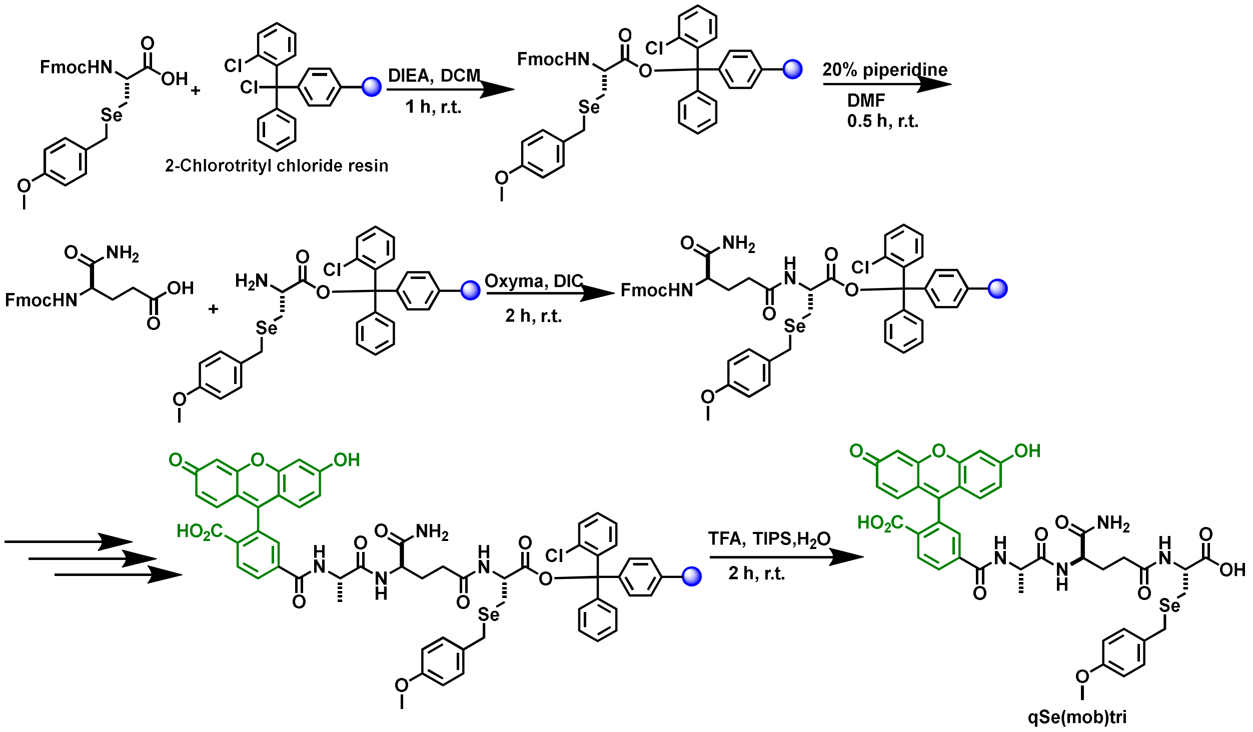


Fmoc-L-Sec(Mob) 1.1 eq was added to a 25 mL peptide synthesis vessel charged with 2-chlorotrityl chloride resin (1.42 g/mol loading capacity) and DIEA (4.4 eq) in approximately 5mL of anhydrous DCM. The resin was shaken for 2 h at ambient temperature and washed with methanol and DCM (3 x 15 mL each). Removal of the Fmoc protecting group was achieved through 20% piperidine in DMF (15mL) for 30 min at ambient temperature, then washed as previously mentioned. Then, 5 eq of Fmoc-D-glutamic acid α-amide were added to the peptide synthesis vessel with 5 eq of ethyl cyanohydroxyiminoacetate (oxymaPure) and 5 eq of N,N'-Diisopropylcarbodiimide (DIC) in DMF (10mL). The resin was shaken at ambient temperature for 2h, then washed as mentioned previously. Fmoc deprotection was done in the same manner as before. Then, 5 eq of Fmoc-L-alanine was added to the resin with 5 eq of oxyma and DIC in DMF (10mL). The resin was shaken for 2h and washed. After Fmoc deprotection of the L-alanine residue, 3 eq of 5(6)-carboxy-fluorescein along with 3 eq HBTU and 6eq DIEA in DMF (10 mL) were added to the resin and was shaken overnight at ambient temperature. The resin was washed as before and added to a solution of TFA/$H_2O$/TIPS (95:2.5:2.5, v/v/v) with shaking for 2 h at ambient temperature. The resin was separated by filtration, and the resulting solution was concentrated *in vacuo* and triturated with cold diethyl ether, affording the isolation of **qSe(Mob)tri**. The peptide was characterized by mass via matrix-assisted laser desorption ionization time-of- flight (MALDI-TOF) mass spectrometry (Shimadzu 8020). The same synthetic path was use to obtain the precursor for qSeDAPtri except 5(6)-carboxyfluorescein was replaced with 5(6)-carboxy- tetramethylrhodamine.

**Scheme S2. Synthesis of qSeDAPtetra precursor qSe(mob)tet**

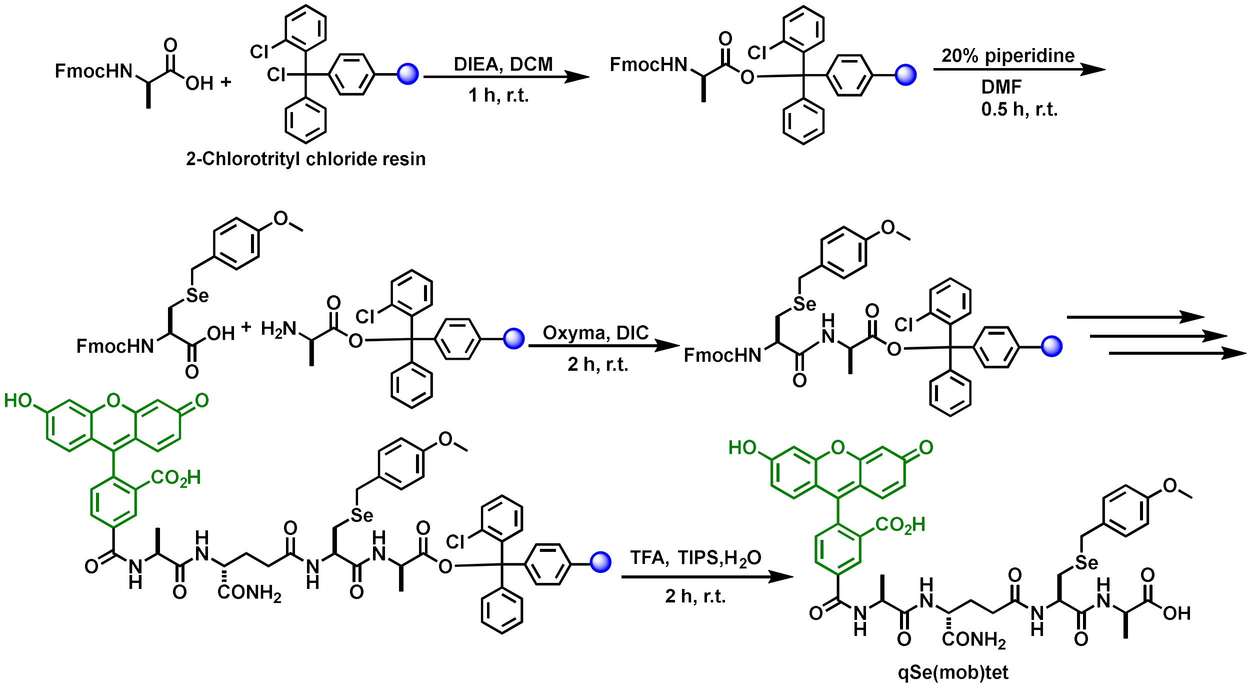

#### Scheme S3: Synthesis of qSeDAPpen precursor qSe(mob)pen

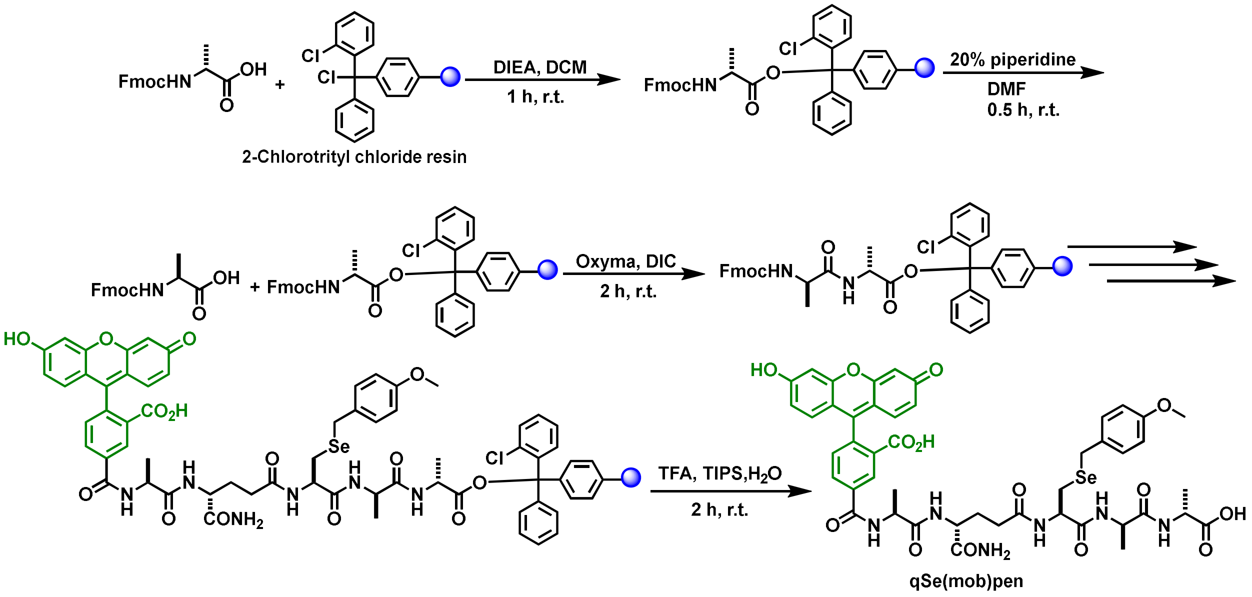

**Scheme S4: General synthetic strategy used to obtain qSeDAPtri, qSeDAPtetra, qSeDAPpen from respective precursors**

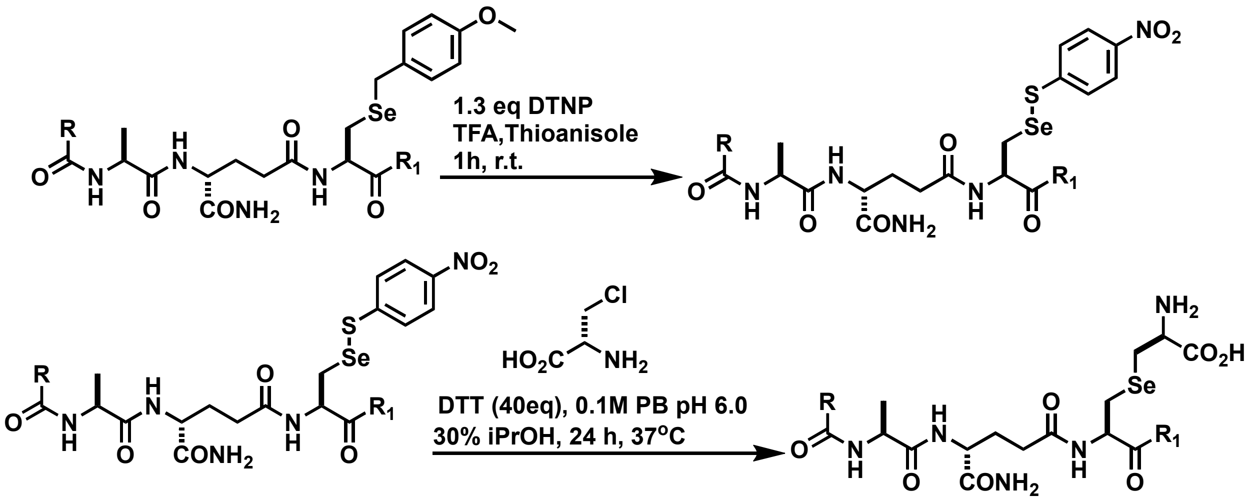


| | R | $R_1$ |
|---|---|---|
| qSeDAPtri | 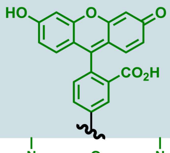 | 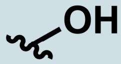 |
| Tm_acceptor | 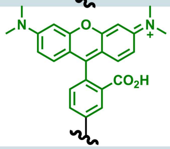 | 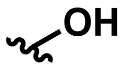 |
| qSeDAPtet | 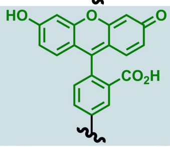 | 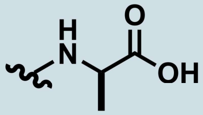 |
| qSeDAPpen | 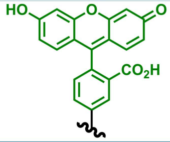 | 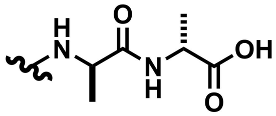 |

Crude **precursor** was treated with TFA:thioanisole (97.5:0.25, v/v) and 1.3 equivalents of 2,2-dithiobis(5-nitropyridine) (DTNP) for 1 h at room temperature to *in situ* swap out the Mob group for TNP. This intermediate was concentrated *in vacuo and* triturated with cold diethyl ether. The TNP peptide was allowed to react with 10 equivalents of β-chloro-D-alanine in 0.1 M phosphate buffer pH 6, and isopropanol (70:30, v/v) containing 40 equivalents of dithiothreitol (DTT) in a round-bottom flask under $N_2$ inert conditions. This reaction proceeded for 24 h at 40 ºC, was quenched with 5% TFA, and isopropanol was removed under reduced pressure by rotary evaporation. The final product was then purified by reverse-phase preparative high-performance liquid chromatography (RP-HPLC) as mentioned above.

**qSeDAPtri** structure & characterization

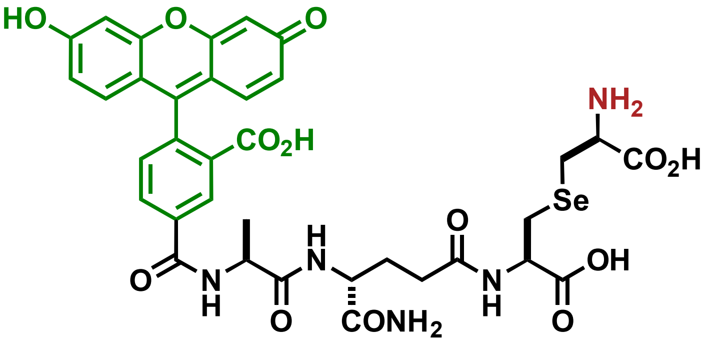


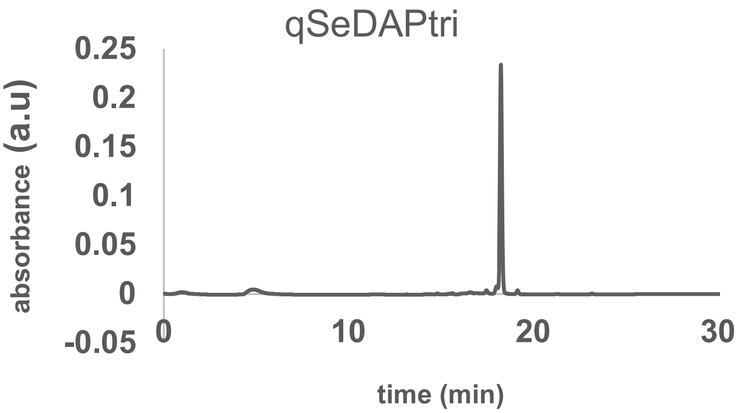


HRMS [M + H$^+$] found: 814.1497

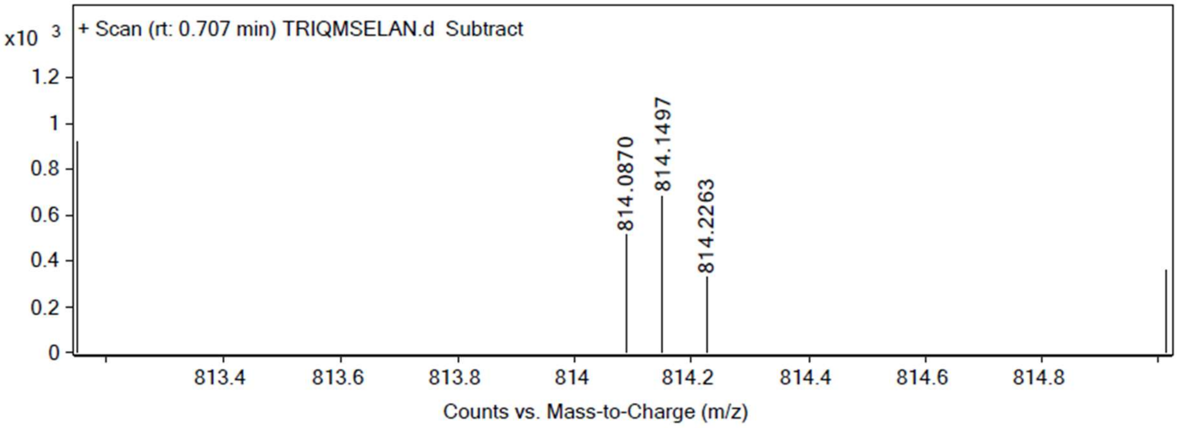

**qSeDAPtri** structure and characterization

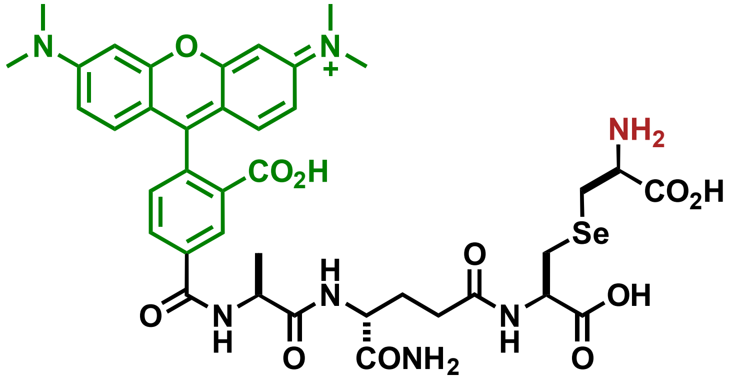


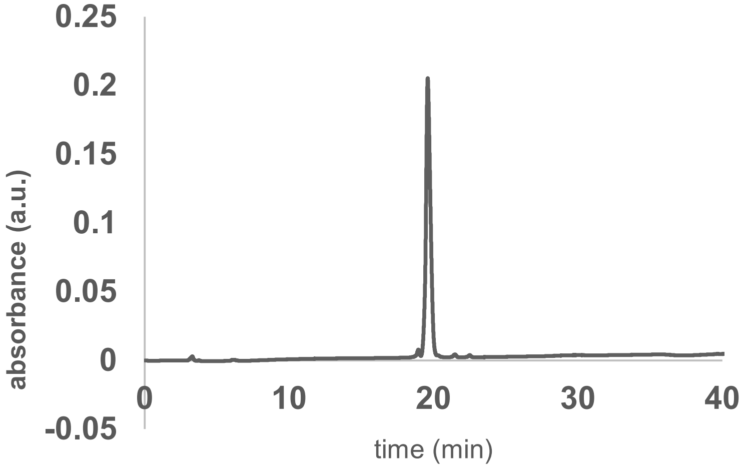


HRMS[M + H⁺] found: 868.2404

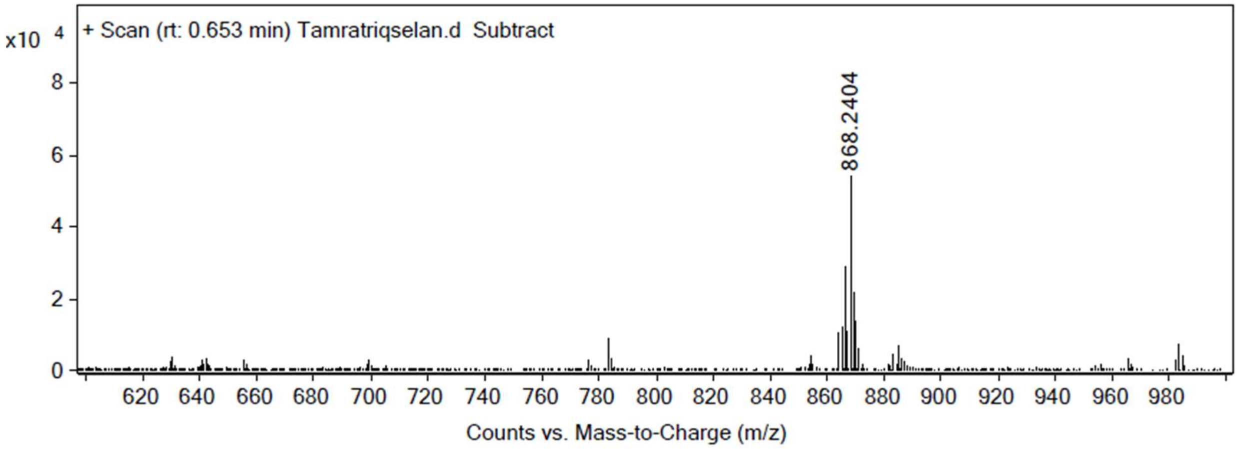

**qSeLYStri** was synthesized as we reported before.[6] Briefly, Crude **qSe(Mob)tri precursor** was treated with TFA:thioanisole (97.5:0.25, v/v) and 1.3 equivalents of 2,2-dithiobis(5-nitropyridine) (DTNP) for 1 h at room temperature to *in situ* swap out the Mob group for TNP. This intermediate was concentrated *in vacuo and* triturated with cold diethyl ether. The TNP peptide was allowed to react with 10 equivalents of 2-chloroethylamine in 0.1 M phosphate buffer, pH 6, and isopropanol (70:30, v/v) containing 40 equivalents of dithiothreitol (DTT) in a round-bottom flask under $N_2$ inert conditions. This reaction proceeded for 24 h at 40 ºC, was quenched with 5% TFA, and isopropanol was removed under reduced pressure by rotary evaporation. The final product was then purified by reverse-phase preparative high-performance liquid chromatography (RP-HPLC) as mentioned above.

Structure & characterization

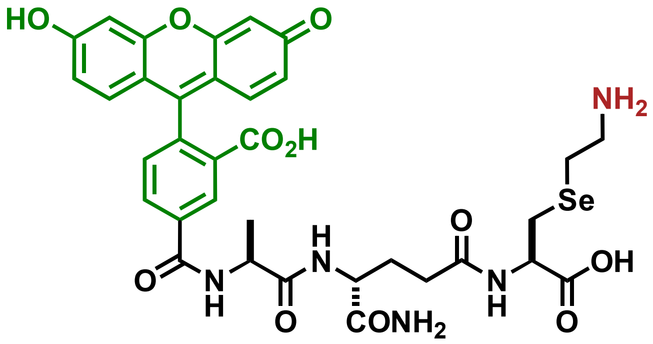


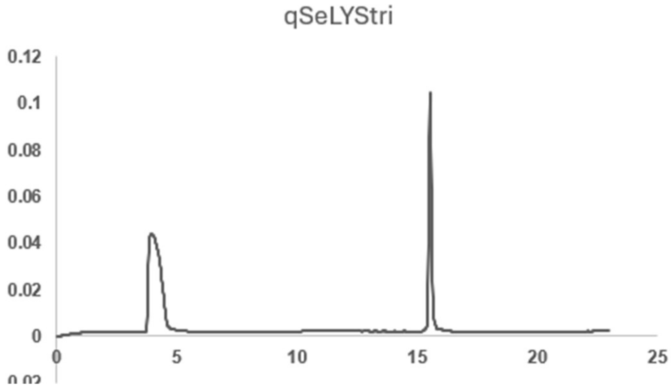


HRMS[M + H$^+$] found: 770.1597

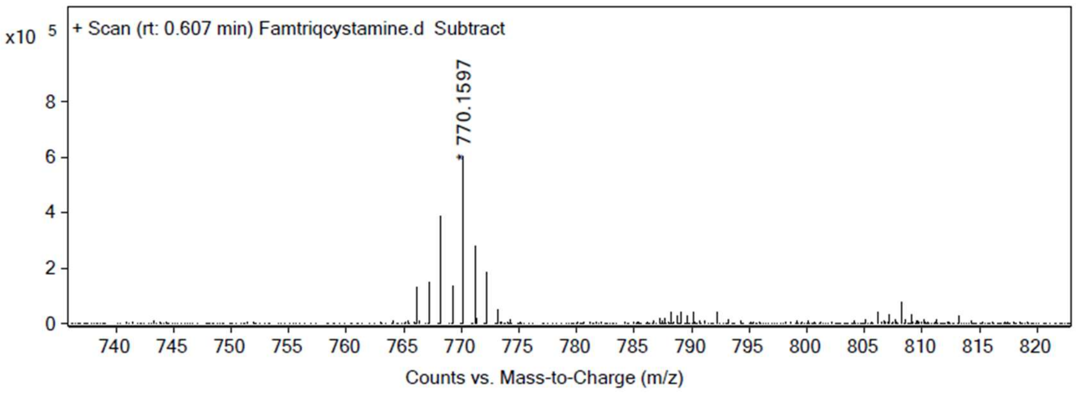
x10 5
+ Scan (rt: 0.607 min) Famtriqcystamine.d Subtract
* 770.1597
0
2
4
6
8
740 745 750 755 760 765 770 775 780 785 790 795 800 805 810 815 820
Counts vs. Mass-to-Charge (m/z)

**qSeDAPtet** structure & characterization

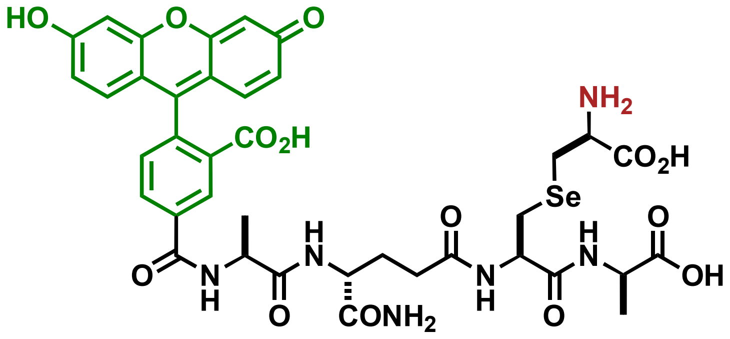


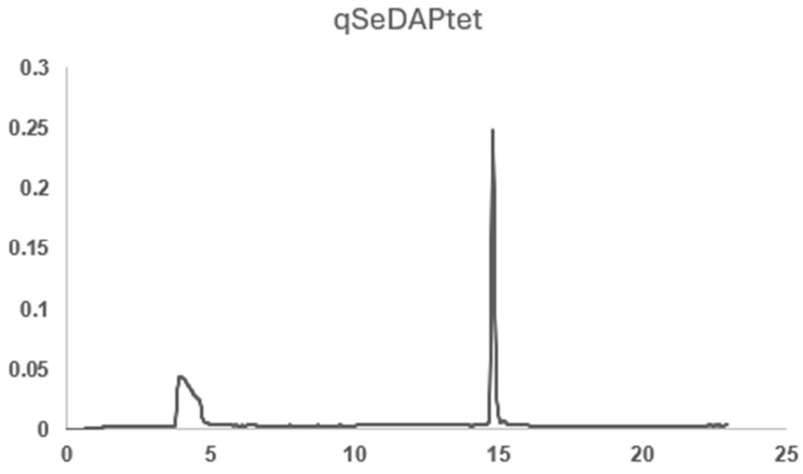


HRMS[M + H+] found: 884.1232

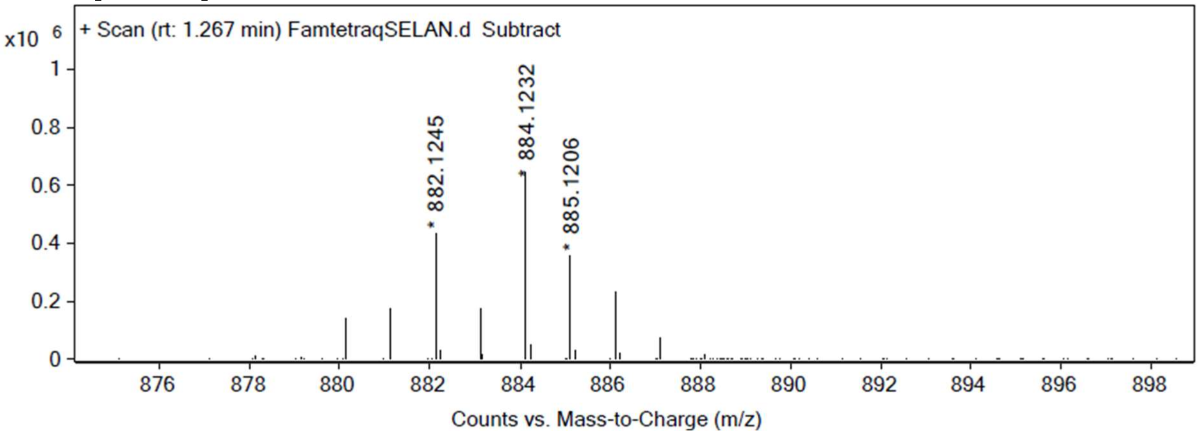

**qSeDAPpen** structure & characterization

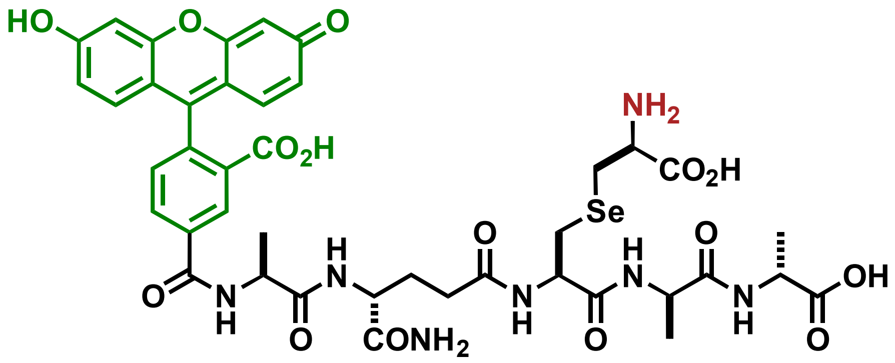


qSeDAPpen

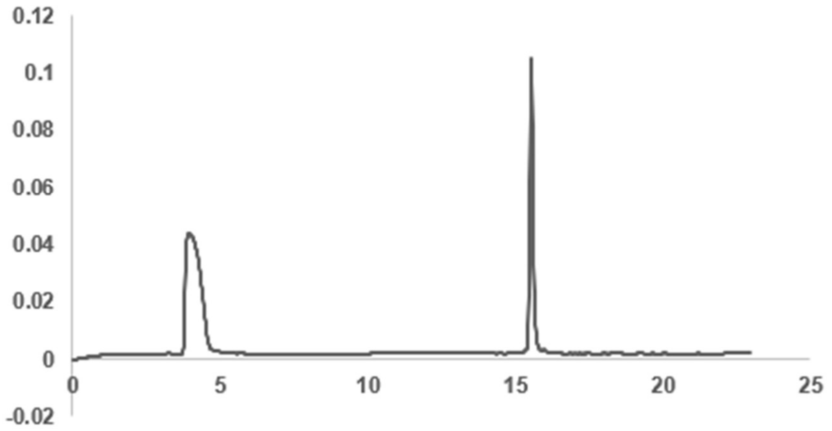


HRMS[M + H$^+$] found: 956.2233

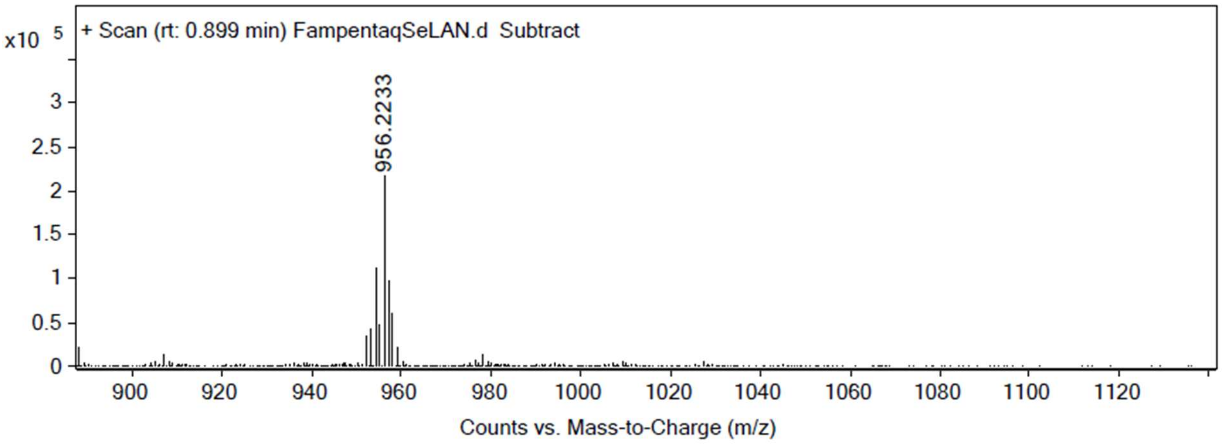

**qCYTtri** was synthesized following previous procedures.[8]

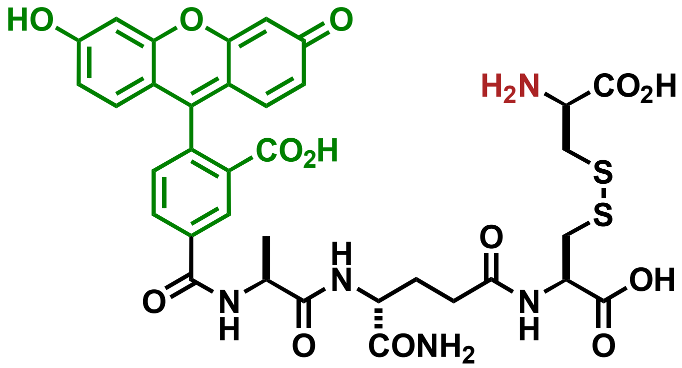


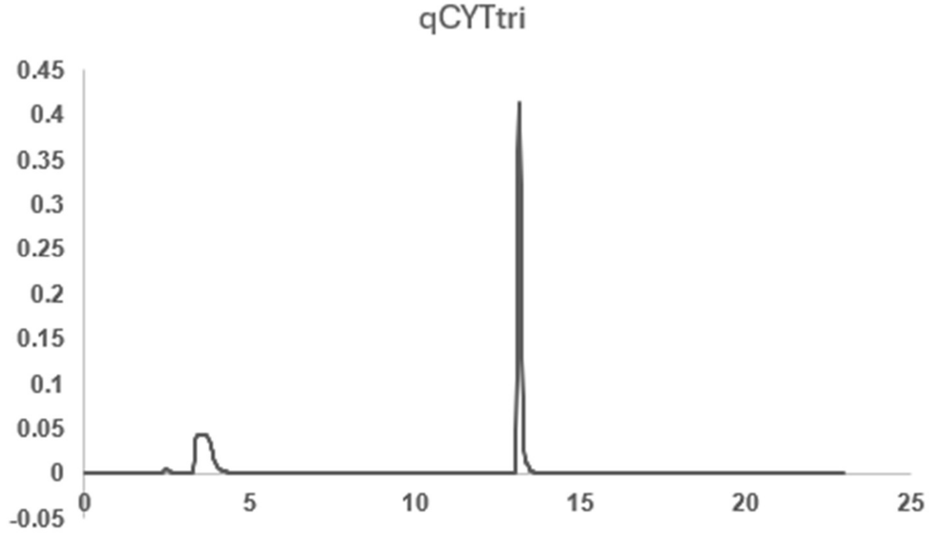


HRMS[M + H+] found: 798.1750.1497

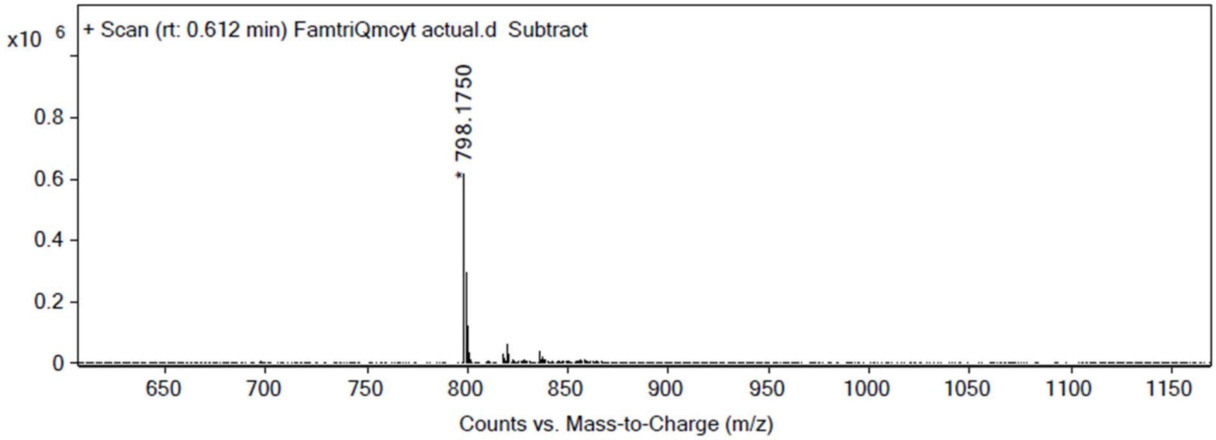

**qLYStri** structure & characterization

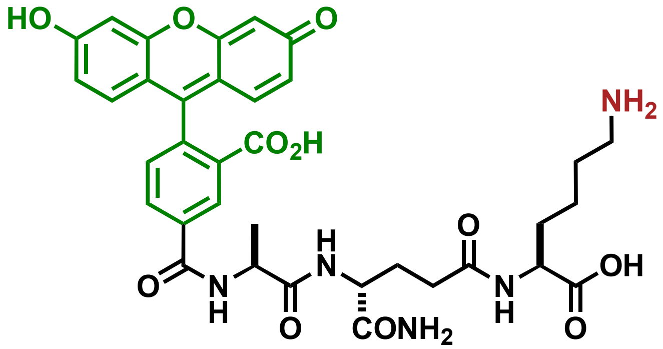


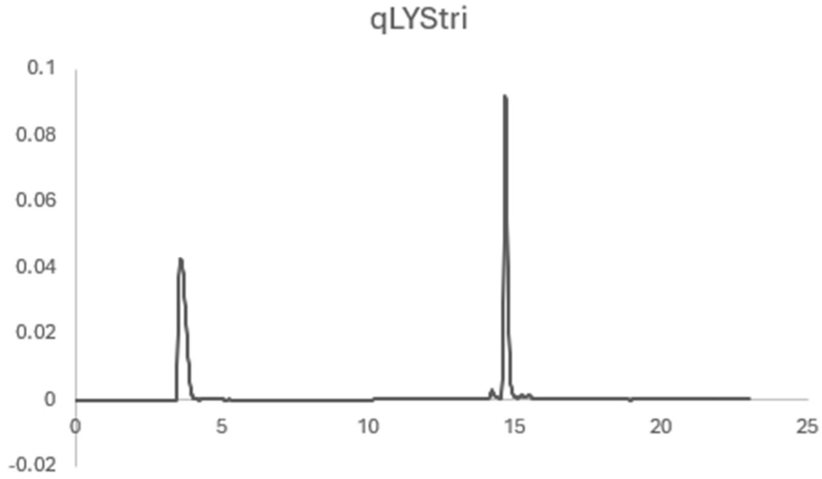


HRMS [M + H$^+$] found: 704.2597

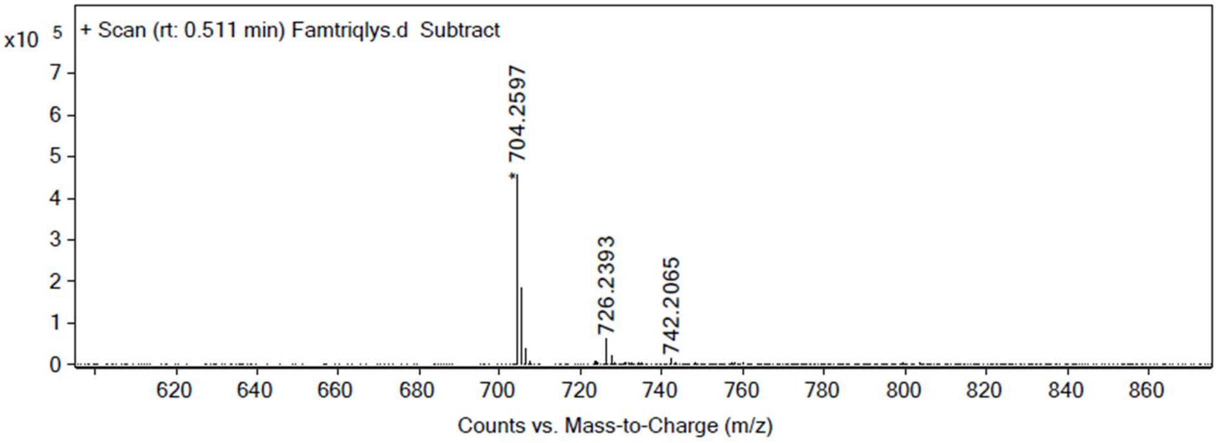

**eLYStri** structure & characterization

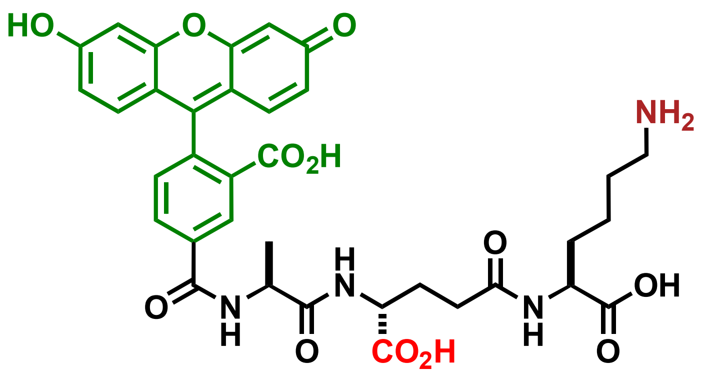


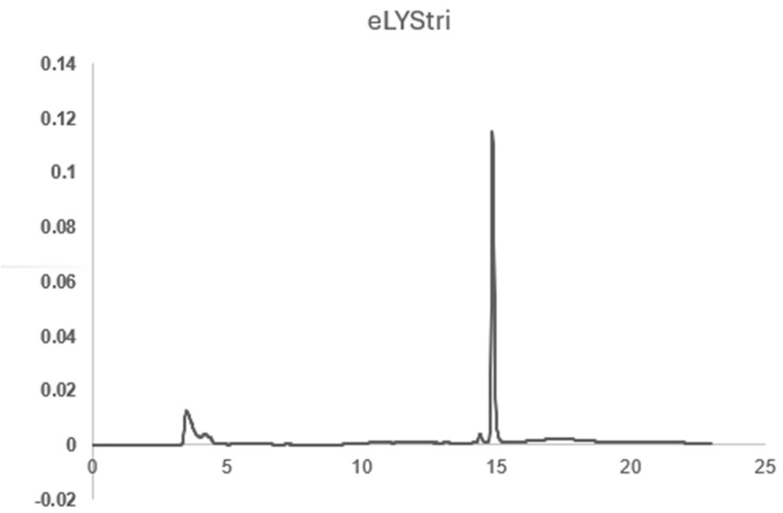


HRMS[M + H$^+$] found: 705.2498

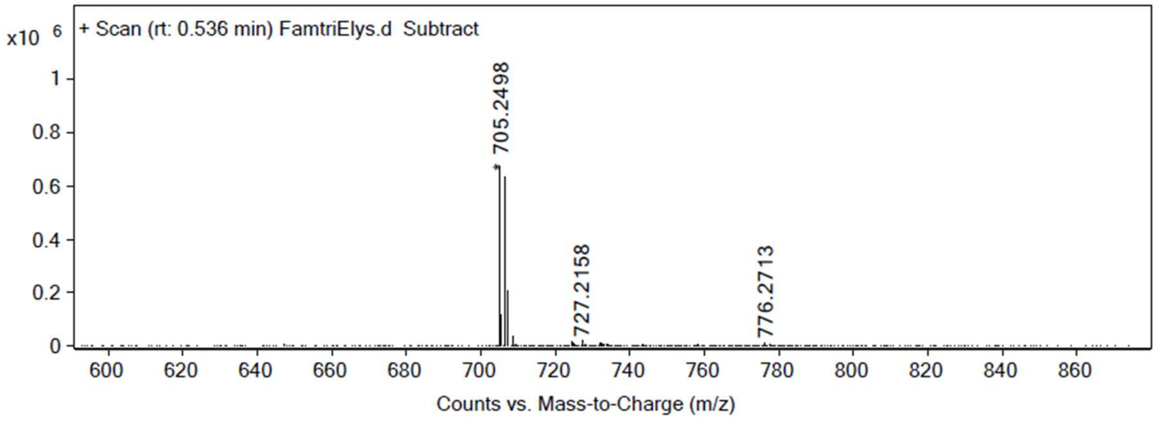

**qLYStet** structure & characterization

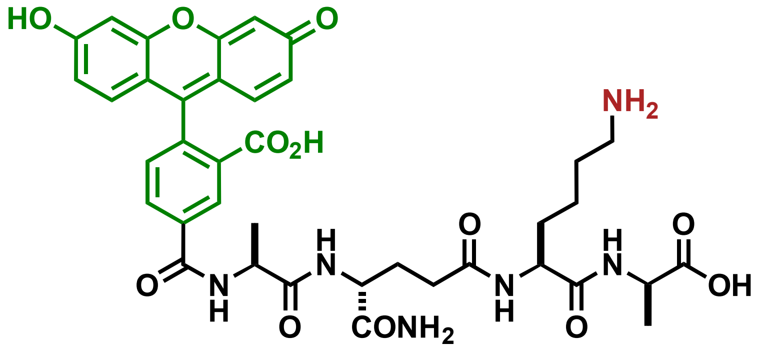


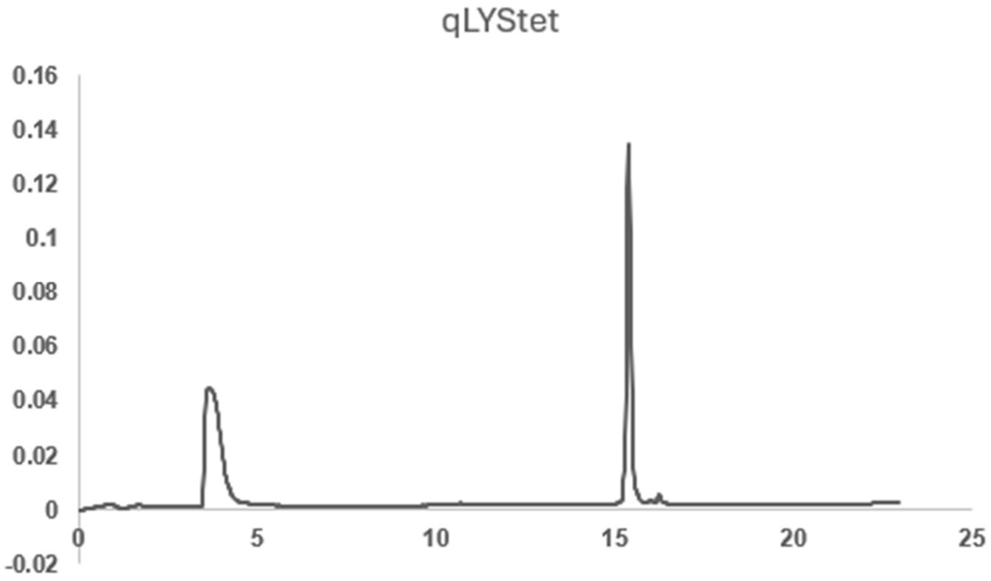


HRMS[M + H$^+$] found: 775.2999

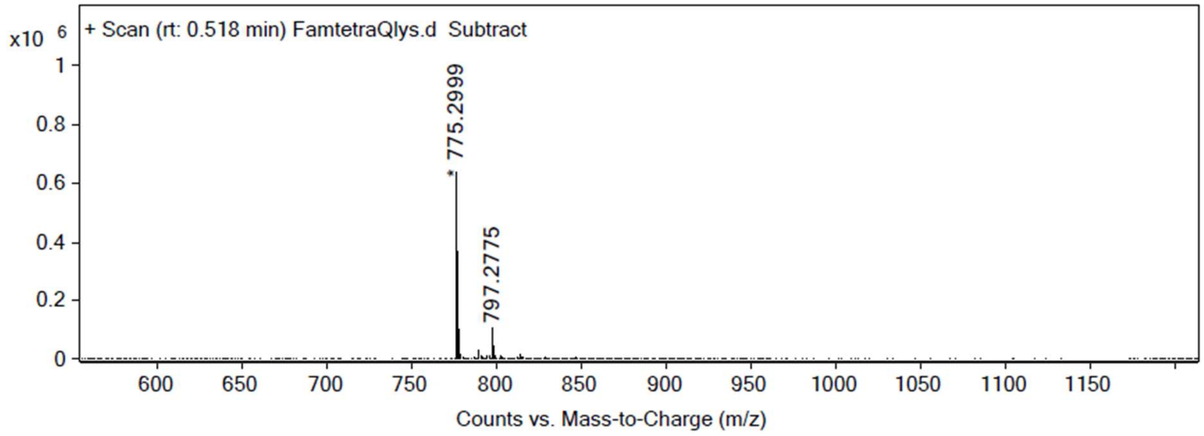

**eLYStet** structure & characterization

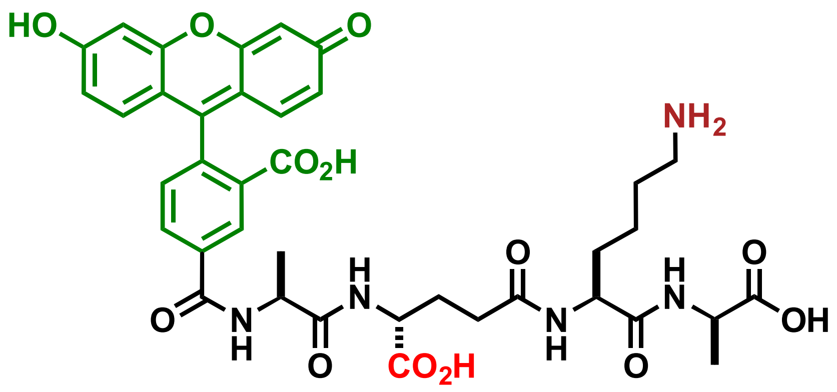


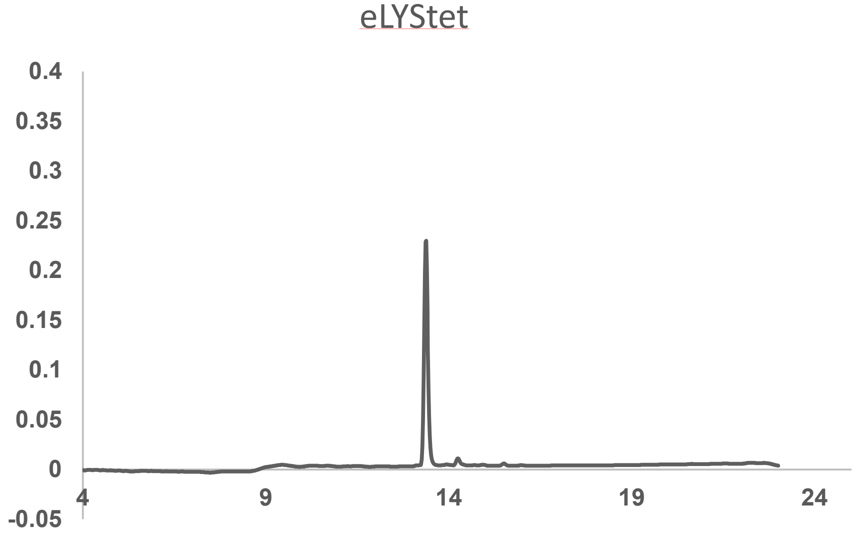


HRMS[M + H$^+$] found: 776.2800

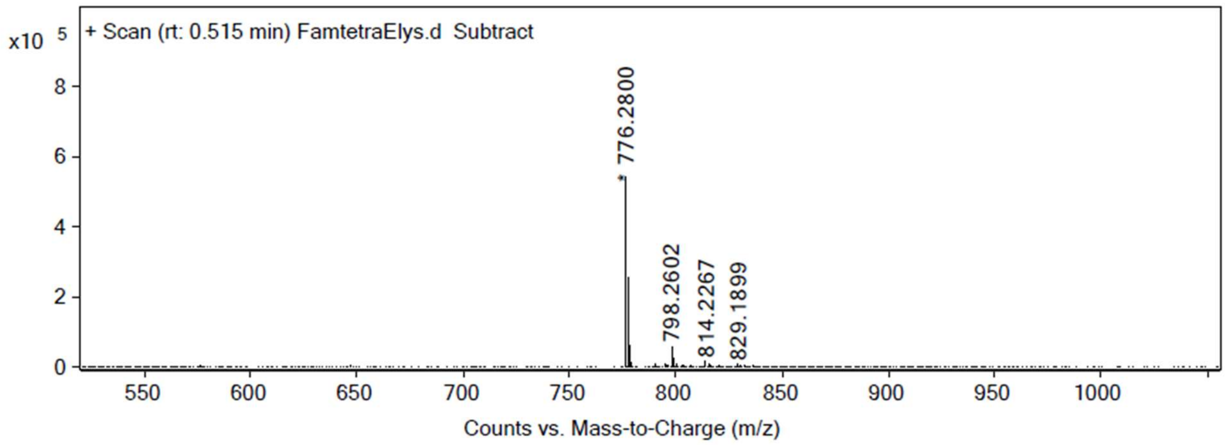

**Scheme S6. Synthesis of tetAcK-yne**

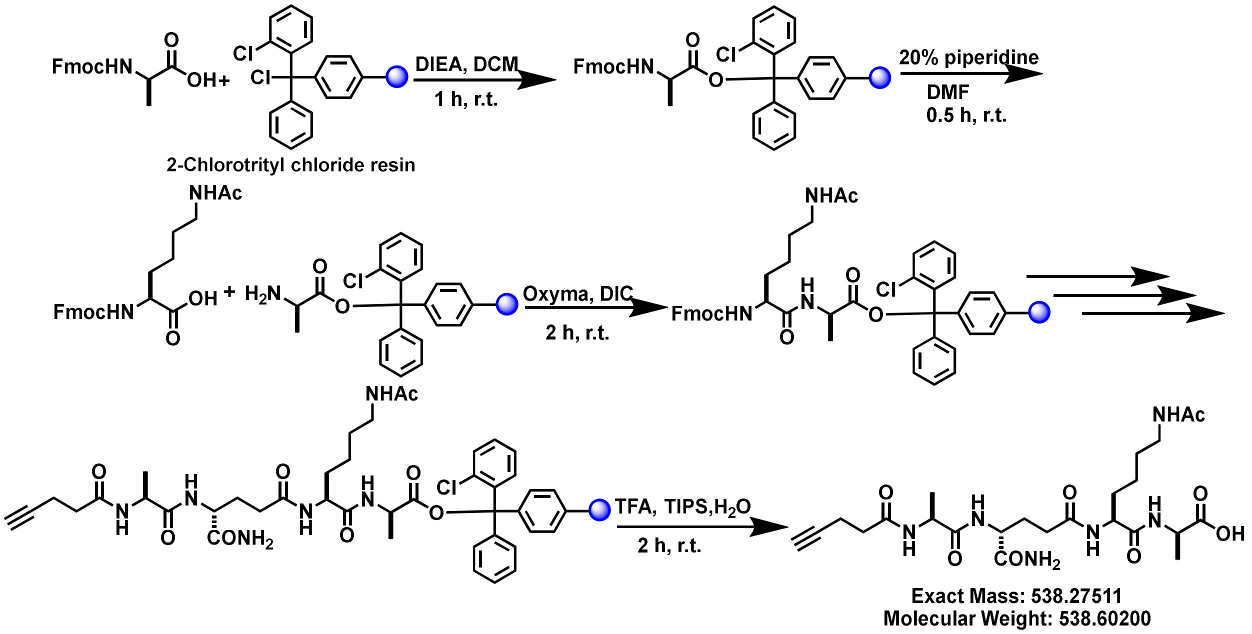


Fmoc-D-Ala-OH 1.1 eq was added to a 25 mL peptide synthesis vessel charged with 2-chlorotrityl chloride resin (1.42 g/mol loading capacity) and DIEA (4.4 eq) in approximately 5mL of anhydrous DCM . The resin was shaken for 2 h at ambient temperature and washed with methanol and DCM (3 x 15 mL each). Removal of the Fmoc protecting group was achieved through 20% piperidine in DMF (15mL) for 30 min at ambient temperature, then washed as previously mentioned. Then, 5 eq of Fmoc-L-Lys(Ac) were added to the peptide synthesis vessel with 5 eq of ethyl cyanohydroxyiminoacetate (oxymaPure) and 5 eq of N,N'-Diisopropylcarbodiimide (DIC) in DMF (10mL). The resin was shaken at ambient temperature for 2h, then washed as mentioned previously. Fmoc deprotection was done in the same manner as before. Then, 5 eq of the remaining amino acid, namely Fmoc-D-glutamic acid α-amide and Fmoc-L-alanine, was added and deprotected in the same fashion. After the Fmoc- deprotection of the final amino acid, 5 eq of 4-pentynoic acid was added to the resin along with 5 eq of oxymaPure and DIC. Following the 2h coupling step, the resin was washed as before and added to a solution of TFA/$H_2O$/TIPS (95:2.5:2.5, v/v) with shaking for 2 h at ambient temperature. The resin was separated by filtration, and the resulting solution was concentrated *in vacuo* and triturated with cold diethyl ether, affording a white pellet of **tetAcK-yne**. The diethyl ether was properly decanted in the waste. The peptide was then purified by reverse-phased preparative high-performance liquid chromatography (RP-HPLC) using waters 1525 pump and 2489 UV/Vis detector and a Phenomenex Luna 10 µm C8(2) 100 Å (250x 21.2 mm) column using a gradient elution with $H_2O$/MeCN with 0.1% TFA at a flow rate of 10mL/min. The HPLC fractions collected were characterized by mass via matrix-assisted laser desorption ionization time-of- flight (MALDI-TOF) mass spectrometry (Shimadzu 8020). The desired fractions corresponding to the purified compound were concentrated by rotary evaporation and then lyophilized using Labconco Freezone 4.5L freeze dryer system. Purity of the peptide was assessed using analytical RP-HPLC using Waters 1525 pump and 2489 detector equipped with a Phenomenex Luna 5µ C8(2) 100Å (250 x 4.60 mm)

column; gradient elution with $H_2O/CH_3CN$ with 0.1% TFA. High resolution mass spectrometry analysis of the peptides synthesized were performed using Agilent 6545 Quadrupole Time-of-Flight (Q-TOF) LC/MS system equipped with Mass hunter workstation and Mass hunter qualitative analysis navigator.

**tetAcK-yne** structure & characterization

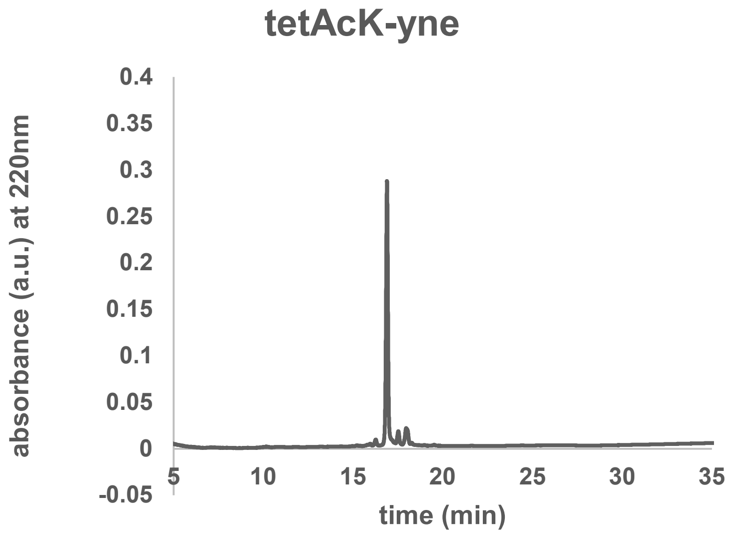


HRMS[M + $H^+$] found: 539.2867

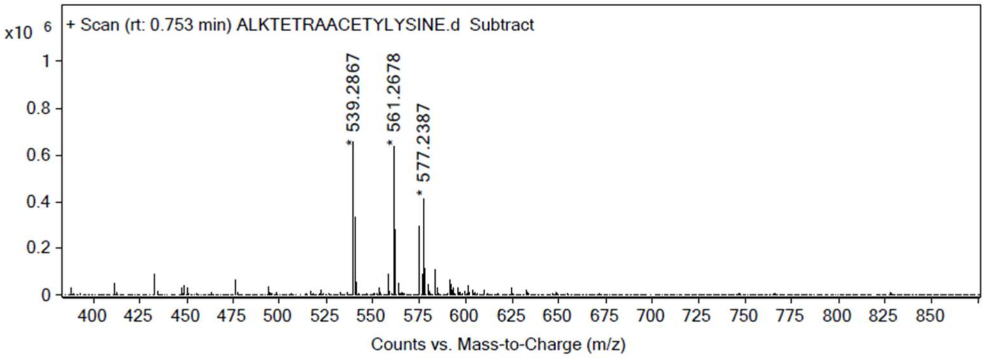
